# Defining the resilience of the human salivary microbiota by a 520 days longitudinal study in confined environment: the Mars500 mission

**DOI:** 10.1101/2020.04.08.031401

**Authors:** Giovanni Bacci, Alessio Mengoni, Giovanni Emiliani, Carolina Chiellini, Edoardo Giovanni Cipriani, Giovanna Bianconi, Francesco Canganella, Renato Fani

## Abstract

The human microbiota plays several roles in health and disease but is often difficult to determine which part is in intimate relationships with the host vs. the occasional presence. During the Mars500 mission, six crewmembers lived completely isolated from the outer world for 520 days following standardized diet regimes. The mission constitutes the first spaceflight simulation to Mars and was a unique experiment to determine, in a longitudinal study design, the composition and importance of the resident vs. a more variable microbiota—the fraction of the human microbiota that changes in time and according to environmental conditions—in humans. Here we report the characterization of the salivary microbiota from 88 samples taken during and after the mission for a total of 720 days. Amplicon sequencing of the V3-V4 region of 16S rRNA gene was performed and results were analyzed monitoring the diversity of the microbiota while evaluating the effect of the three main variables present in the experimental system: time, diet, and individuality of each subject. Results, though showing statistically significant effects of all three variables, highlighted a main contribution of salivary microbiota personalized features, that is an individual-based resilience of the microbiota. Such findings open the way to consider salivary microbiota under the light of a pronounced personalization even after sharing the same physical space for more than a year.

## Introduction

The host-associated microbiota is stirring the attention from many fields of life science, including basic biology, evolutionary studies, biomedicine, and biotechnology. It is now well known that it plays several roles in modulating the host health and that changes in the composition of the microbiota in specific human body districts or organs (e.g. skin, gut, vagina, lung) may influence the correct functionality of other organs.^1^ The concept of holobiont reflects the intimate relationships between the host and the microbiota^2,3^ but it is often difficult to determine which part of the host-associated microbiota is in intimate relationships with the host vs. an occasional presence. Cross-sectional studies have been used to decipher the more stable, core, microbiota, present in all individuals analyzed, in comparison with the fraction which is more variable, i.e. present in few individuals only (see for instance^4^) and longitudinal analyses helped to understand the temporal stability of the microbiota.^5^

The human microbiota is not a single entity but it may have different characteristics and roles. The gut microbiota is expected to be more stable over time than other cavities that are more exposed to the environment—the oral cavity represents one of the first entry point of our body and is thus massively influenced by environmental conditions. The salivary microbiota is known to be affected by both biotic and abiotic factors,^6,7^ including age, saliva chemical composition, tongue, and teeth.^8^ Consequently, it is still under debate how much of the oral microbiota is stable over time and if this stability can be considered as a tight association with the host.^7,9,10^ Given its sensibility to external perturbations, the salivary microbiota could be a good model to inspect the temporal dynamics and subject-by-subject variations impacting the human microbiota, but this sensibility could be a double-edged sword. Even if the disclosure of salivary microbiota temporal stability, and/or subject individuality, could indeed impact on scientific fields spanning from personalized medicine to forensic microbiology, controlling environmental exposures of salivary microbiota is difficult especially during our every day life. Standardize these perturbations implies isolation procedures that are difficult to put in place.

Mars500 was the first long-term international study into interplanetary space flights. Managed by the European Space Agency and the Russian Space Agency, it was conducted in 2010-2011 when six male volunteers were kept for 520 days in a common confined environment established by the Institute of Biomedical Problems (IBMP) in Moscow, simulating a space flight to Mars. Data from Mars500 mission were studied from various point of view, including behavior,^11^ effect of cultural background,^12^ cognitive performances,^13^ circadian rithms,^14^ hormone levels,^15^ and surface and gut microbiota.^16,17^ Mars500 hence constitutes a unique experiment to determine, in a longitudinal study design, the composition and importance of the resident microbiota vs. a more variable microbiota (changing with time and environmental conditions) in humans. Aim of the work was to inspect the temporal dynamics of salivary microbiota, assessing the effect of diet regimes and individuality, using Mars500 as a unique long-term experiment where subjects were all confined in the same shared environment.

## Results

### Salivary microbiota composition during the study

To inspect how the salivary microbiota reacts in a confined environment, we characterized samples collected during the entire duration of the Mars500 mission (720 days in total) by 16S rRNA gene amplicon sequencing of the variable region V3-V4. Table S1 summarizes all phases of the mission whereas Figure S1 reports the sampling scheme used in this work. Amplified sequences formed 1890 amplicon sequence variants (ASVs) with a median number of 172.00 ASVs per sample (ranging from 81 to 317). A total of 4,337,540 sequences specifically aligned to an ASV resulting in a sequencing depth ranging from 20,084 to 116,809 and a median value of 47,044 (for additional information about sequence analysis pipeline and the number of sequence obtained in each pre-processing step see Supplementary material, Supplementary Table S2, and Supplementary Figure S2). All replicates reported an accuracy higher than 0.96 with a Spearman’s rank correlation (*ρ*) that ranged between 0.94 and 0.98 (Supplementary Table S3, S4 and Supplementary Figure S3). Rarefaction curves reached a plateau above 15k reads suggesting an adequate sequencing depth for all samples (Supplementary Figure S4). Good’s coverage estimator ranged between 99.99% and 100.00% across all samples indicating that roughly 0.01% of the reads in a sample came from ASVs that appear only once in that sample (Supplementary Table S5).

Roughly, 99% of sequences aligned to variants that came from known bacterial taxa (Table S2). Supplementary Table S6 shows the overall taxonomic composition of samples whereas Figure S5 reports the phylogenetic tree reconstructed from ASVs. At phylum level Firmicutes, Bacteroidetes, Actinobacteria, Proteobacteria, and Fusobacteria accounted for more than 97% of the total number of reads assigned to taxonomically annotated ASVs (Figure 1a and Supplementary Table S6). The total bacterial diversity (namely the alpha diversity) remained constant during the mission with no significant differences detected between the isolation period and the follow-up, across different diets, and across subjects (Table S7 and Figure 1 panels b, c, and d). Also time did not impact bacterial diversity as showed in Figure 1 (random mixed model fitted using crewmembers as random intercept: slope lower than 0.002; Supplementary Table S8).

**Figure 1:**
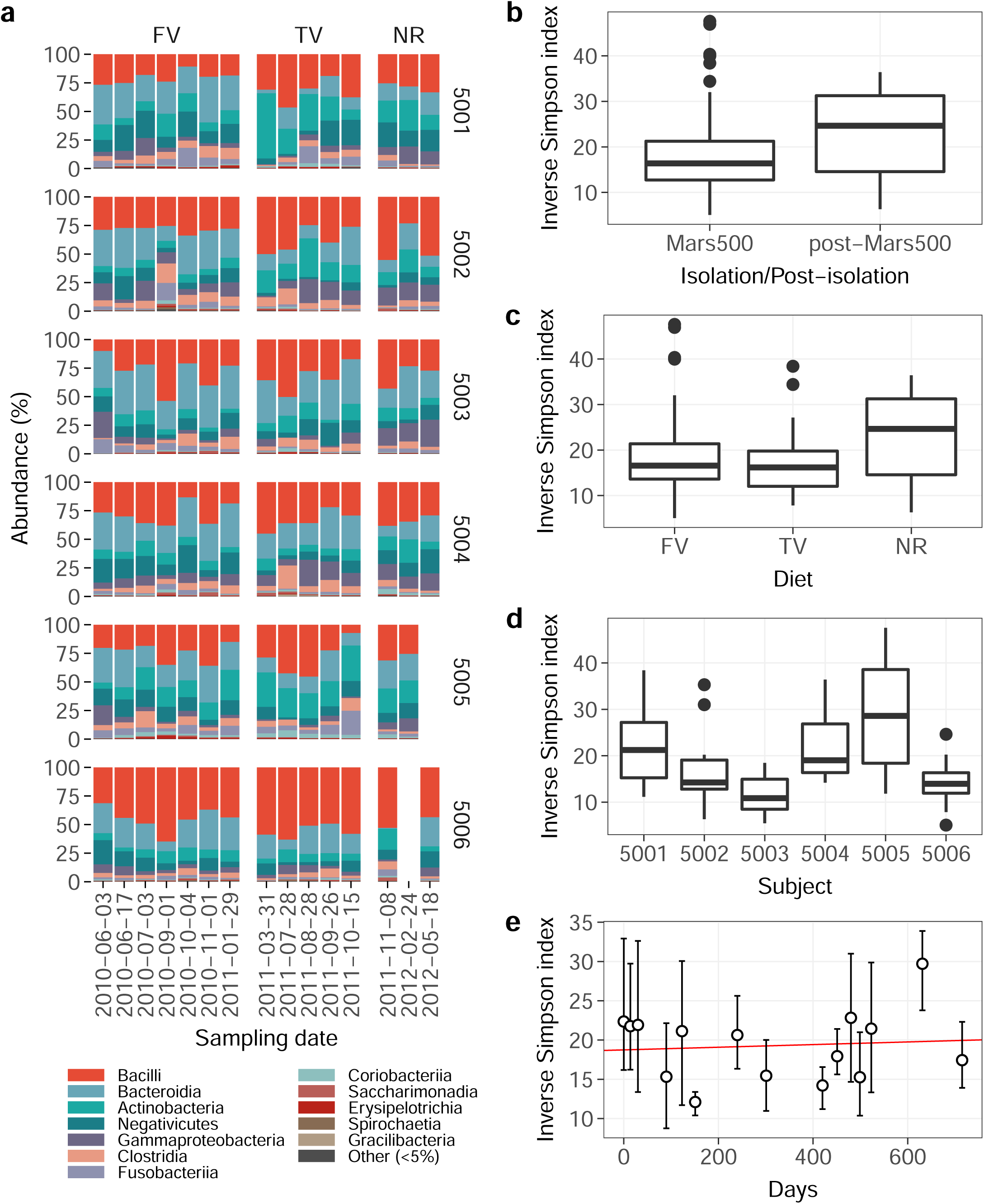
Salivary microbiota diversity along Mars500 mission and follow-up. a) Distribution of the main bacterial classes. Panels were divided according to crewmembers (vertically) and diets (horizontally). ASVs with a relative cumulative frequency lower than 5% in all samples were collapsed into a single group called “Other”. b) Differences in alpha diversity— reported using the inverse Simpson index just like panels c, d, and e—between samples collected during and after the isolation period. c) Differences across diets (FV, first variant; TV, third variant; NR, normal diet). d) Differences among subjects. e) Differences along the whole mission and during the follow-up. Points are the average diversity values among subjects whereas errorbars represent the 95% confidence interval around the mean. The red line represents the population effect of the linear mixed model whose coefficients are reported in Table S8.

### Effect of food and time on salivary microbiota

We inspected differences across samples (namely beta diversity) using non-metric multidimensional scaling (nMDS) on quantitative and qualitative indexes. Samples showed a similar distribution with all index tested (Figure 2a): Sorensen index and unweighted unifrac distance (qualitative analysis), and Bray-Curtis and weighted unifrac distance (quantitative analysis). As opposed to alpha diversity, subjects, diets, and time significantly contributed to shape the salivary microbiota with different percentage of variance explained depending on the index but never exceeding 10% of the total variance (Figure 2b). For all diversity indexes (except for the Sorensen index which reported a significant effect of subjects) the dispersion of tested factors was homogeneous, meaning that only the composition of samples varied among groups as highlighted by the permutational analysis of variance reported above. Diet impacted on bacterial genera usually present in the salivary microbiota of healty subjects—such as *Actinomyces, Veilonella*, and *Fusobacterium*^18^—but also on *Peptostreptococcus, Haemophilus, Megasphaera*, and Prevotella, which have been correlated to different disorders of the oral cavity (such as periodontitis, dental caries, and oral lichen planus).^19–21^ Bacterial species classified as *Alloprevotella, Fusobacterium, Dialister, Veilonella*, and *Megasphaera* followed the same pattern: decreasing abundance passing from the the first to the third diet and then back to starting values during the follow up (even if the shift was not significant). Other species like *Haemophilus* and *Prevotella* reported a significantly higher abundance during a single diet—namely *Haemophilus* was more abundant during the follow up and the abundance of *Prevotella* was higher during the first diet. At phylum level, diets impacted more on Bacteroidetes and Firmicutes. Whithin them, 4 out of 28 (14%), and 3 out of 7 (43%) genera reported (at least) a significant difference during diet changes (Figure S6).

**Figure 2:**
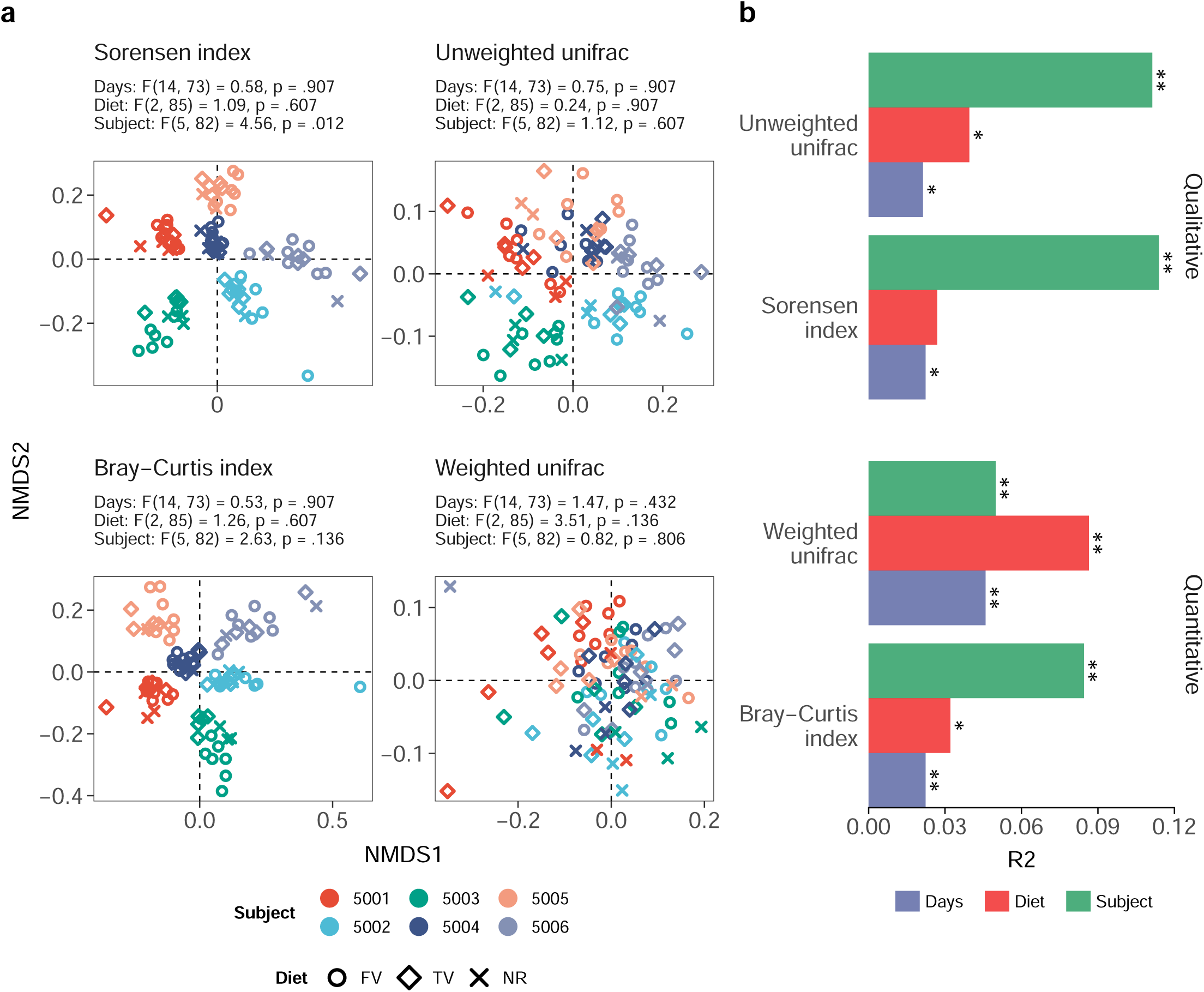
Microbial assemblage variation according to diet, crewmembers, and time. a) Non-metric multidimensional scaling based on different beta-diversity indexes (reported on the top of each panel). Samples were colored according to crewmembers whereas the point shape represents the type of diet (FV, first variant; TV, third variant; NR, normal diet). The dispersion of groups was tested for homogeneity and results were reported on the top of each oridnation (a p-value higher than 0.05 means that dispersions are homogeneous). b) Permutational multivariate analysis of variance using distance matrices based on the same indexes reported in panel a. The R^2^ values associated with each factor used in the analysis is reported in the horizontal axis whereas asterisks report the significance level of each factor (*, p-value < 0.05; **, p-value < 0.01). Colors represent the different factors modeled in the analysis. For additional information about diets, sampling point and crewmembers see Supplementary information and Supplementary Figure S1.

To explore the effect of time on bacterial diversity we used change-point analysis on both within-subject (Figure S7 panel a) and between-subjects diversity (Figure S7 panel b). Within-subject diversity measures changes in the salivary microbiota of each crewmember through time, whereas between-subjects diversity compares the salivary microbiota of different crewmembers at each time point (Figure 3a and b). Three segments significantly divided within-subject diversity with two change-points at 123 days and 480 days. Between-subjects diversity was not segmented since the overall model gave better results than the segmented one according to the genetic algorithm used during optimization (Figure 3b). The overall between-subjects model had an effect size of −0.00004 which means that after 520 days of isolation the overall diversity decreased by 0.02179. The effect of time on within-subject diversity was indeed higher than the one observed for between-subjects diversity. During the first 123 days the effect modeled was 0.00103 reflecting an average increase of 0.12636 for all crewmembers. After the first change-point, within-subject diversity started to decrease with a regression parameter of −0.00064 (average decrease during the second segment of −0.22835). After the second change point, which roughly matched the end of the isolation period (Figure 3c and d), the within-samples diversity started to increase again. At the end of the follow-up period diversity increased again of 0.29474 exceeding the average value detected in the first day of isolation.

**Figure 3:**
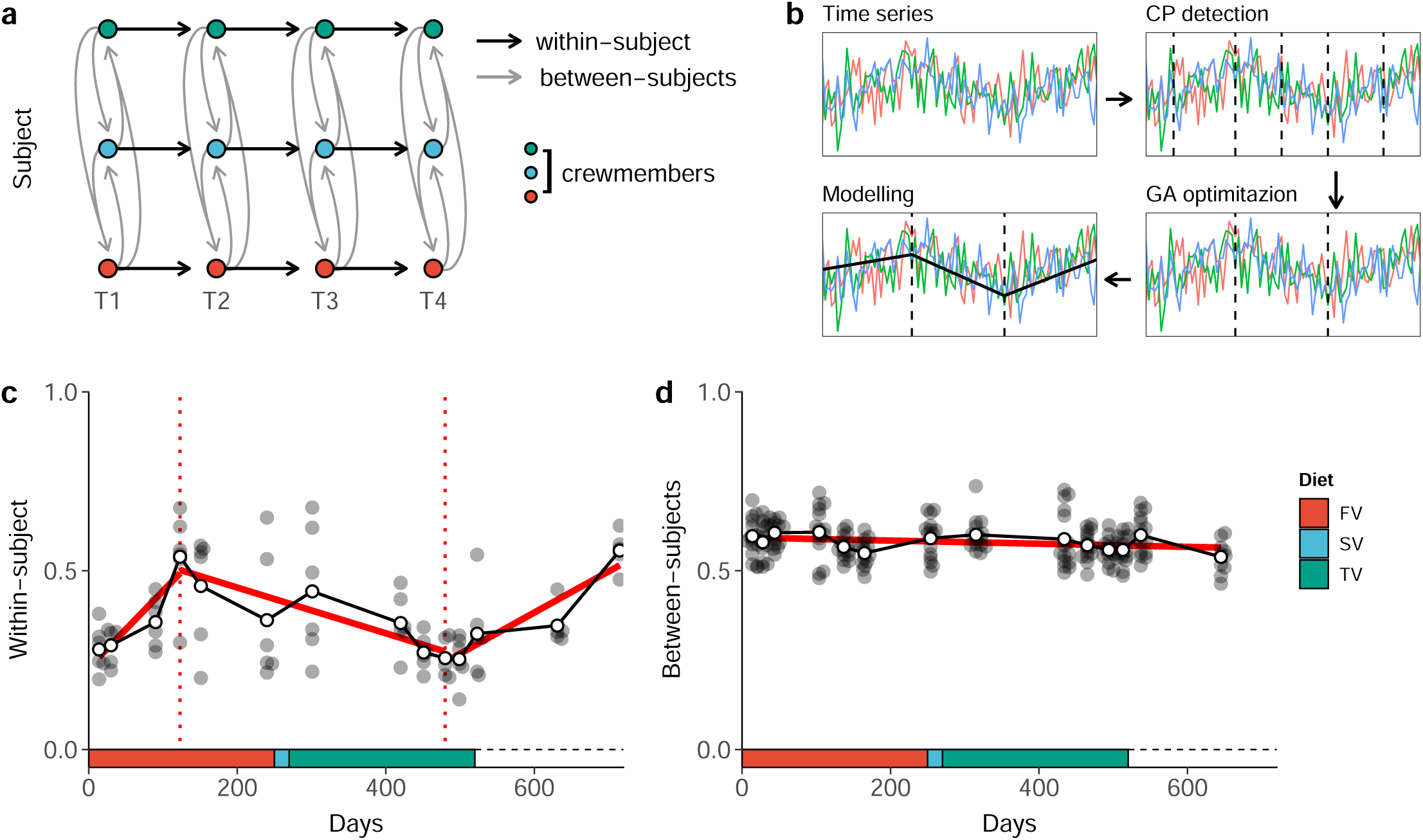
Crewmembers’ salivary microbiota composition in time. a) Bray-Curtis (also known as quantitative Sorensen) index has been used to inspect distances between and within-subject during isolation and follow-up. Between-subjects diversity was computed by comparing the salivary microbiota of each subjects at each timepoint (gray arrows); within-subject diversity was computed by comparing the salivary microbiota of the same subject over time (black arrows). b) Change-point analysis revealed changes in salivary microbiota composition of each subject (CP detection). Genetic algorithm and linear modelling detected increasing/decreasing patterns along time (GA optimization). Finally, we fit a linear mixed-model for each segment detected using crewmembers as random intercept (Modelling). c and d) Results obtained following the pipeline reported in “b” for within- and between-samples differences. Diets were reported in the bottom part of the plots using different colors (FV, fist food variant; SV, second food variant; TV, third food variant). Since crewmemebrs ate freely during the follow-up no diet was reported in the plot.

### Resilience of salivary microbiota

The average abundance of ASVs correlates with their persistence, the number of subjects in which a given ASV was detected at each time point. Figure 4a shows the increasing trend of log-transformed abundance with an R-squared value of 0.72 (*b* = 10.09, 95% CI 9.81, 10.38]). Time-resolved clustering produced two groups of ASVs: one, called inconsistent micriobiome (Cluster 1), included variants detected in a small number of subject at each time point, whereas the other (Cluster 2), called stable microbiota, included variants detected in the vast majority of subjects during the whole mission (Figure 4a and Figure S8, panel a and b). The inconsistent microbiota showed low average persistence in respect with the stable microbiota but it contained the largest amount of variants (1746 ASVs against 144 of stable microbiota). Unlike stable ASVs, subjects lost and acquired inconsistent ASVs both during and after the isolation period (Figure S8 panel b and c). Stable ASVs were detected in roughly 30% of all subjects at each time point (26 samples on 88) with sporadic losses and acquisitions (Figure 4a and Figure S9).

**Figure 4:**
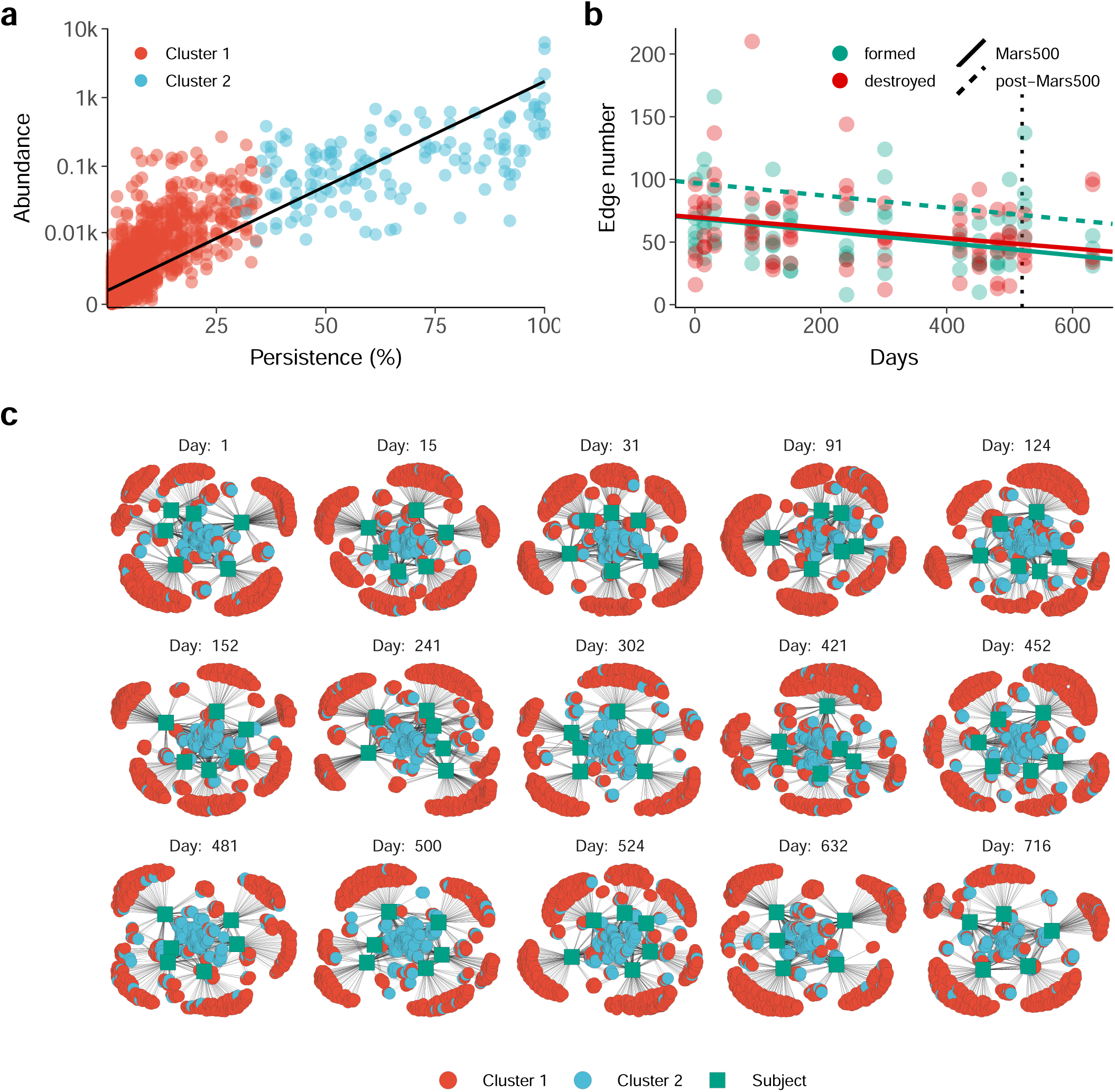
Bacterial community structure in time. a) Persistence and abundance of ASVs detected. Persistence was expressed as the number of subject in which an ASV was detected whereas abundance was expressed as log-normalized number of reads assigned to that ASV (the black line represent results of the linear fitting: 95% CI [9.81, 10.38], t(1926) = 70.27, p < 0.001). Cluster 1 (reported in red) was composed of ASVs with a lower persistence and abundance than Cluster 2 (reported in blue). b) Number of edges formed (green) and destroyed (red) at each time point. Lines represent the result of two linear mixed models with subject as random intercept (Table S9). The dashed line represents the effect at the end of the isolation—it was reported only for formed edges since the number of destroyed edges was not significantly impacted by the isolation. c) Community networks at each time point. Days are reported at the top of each network whereas nodes with no edges (namely ASVs not detected at a given time point) were not reported to save space for graphical representation.

We represented the acquisition and loss of bacterial species during the whole mission using networks. At each time point we linked subjects to ASVs detected in their salivary microbiota forming a bipartite network structure which reflected the underlying bacterial community structure. The loss and acquisition of bacterial ASVs was shown in supplementary video S1 where green squares represent subjects, red circles represent inconsistent microbiota, and light blue circles represent stable microbiota As shown in the video, the topology of the networks did not change in time, but at each time point subjects acquire/release bacterial species from/into the environment, except for stable ASVs which are shared by most crewmembers and thus (almost) always present in central part of the network. The number of new edges formed and destroyed passing from one time point to another slightly decreased in time (mixed effect model 95% CI for formed edges [−0.08, −0.02] and destroyed edges [−0.08, −0.01] Table S9). The end of the isolation period significantly increased the average number of formed edges (namely acquired ASVs) of 28 but the trend was still negative (Figure 4b). The number of formed edges was independent from the number of lost edges (Spearman’s *ρ* = −0.04; p-value = 0.750). The salivary microbiota structure did not change both during and after the isolation period reporting a similar network topology even if we exclude lost ASVs (Figure 4c).

### Drivers of diversity

To inspect drivers of beta diversity along and after the isolation period, we fitted a linear model for each ASV detected in the salivary microbiota of crewmembers. The Mars500 mission time-scale was divided into three stages according to the changepoints detected for within-subject diversity. Crewmembers showed a different number of ASVs reporting a trend of diversity similar to the one reported in Figure 3c. The number of ASVs showing a significant effect of time and changepoints ranged from 3 to 28 depending on the subject (Figure 5a and Table S10). Stable ASVs—namely those grouped into Cluster 2 according to their prevalence in time—that showed a significant trend of diversity enriched the saliva of four out of six crewmembers, if compared with the overall occurrence (Figure 5b). The fraction of stable ASVs in subjects 5004 and 5005 was roughly six times higher than the average fraction of stable ASVs, whereas subjects 5002 and 5006 reported a fraction three times higher than the average.

**Figure 5:**
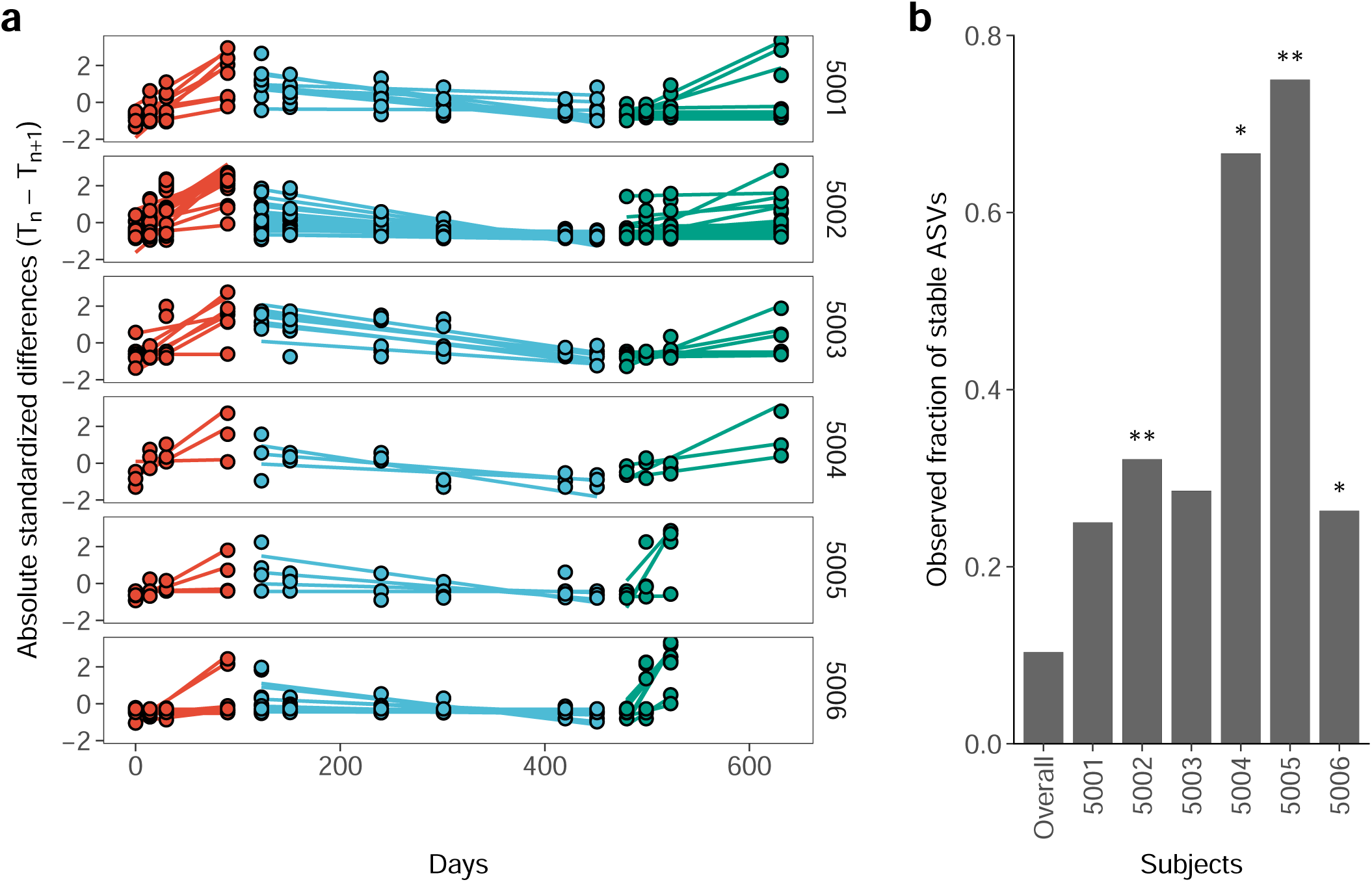
Drivers of diversity for each crewmember. a) Single ASVs showing a trend of diversity along time similar to the whole within-subject diversity were reported. Points represent absolute differences between two consecutive time points whereas colors report the segmentation detected through change-point analysis (Figure 3c). Lines show the predicted differences using linear modelling. We standardize differences to represent ASVs with different ranges of values in the same panel. b) Enrichment analysis of memebers of Cluster 2 in respect with the population. The firs bar on the left represents the overall fraction of AVSs assigned to Cluster 2 whereas the other bars report the fraction of ASVs ssigned to the same cluster for each subject. Adjusted p-values were reported using a single asterisk (p < 0.05) or two (p < 0.01).

## Discussion

The reported experimental results under Mars500 mission evaluated the temporal dynamics of the human salivary microbiota in a controlled and confined environment. All samples from the 6 (male) crewmembers included Firmicutes, Bacteroidetes, Actinobacteria, Proteobacteria, and Fusobacteria as main phyla and conserved their taxonomic composition along time and among individuals, as previously reported by other studies (see for instance^7,9,22,23^). Despite this conservation, external factors significantly influenced the salivary microbiota composition but their influence was restricted to a low percentage of the community (less than 10% of total variance explained). We found that a great number of species intermittently passed through the salivary microbiota but they neither affected its overall structure nor its taxonomic composition. Despite their number, transient species struggle to thrive in human saliva reporting a low abundance during the whole experiment. On the other hand, a hundred bacterial taxa dominated the salivary microbiota composition of all crewmembers rarely changing host and abundance even when isolation period ended. The isolation time affected bacterial diversity of single individuals but it did not alter the microbiota of the whole crew. The bacterial diversity of crewmembers decreased—if compared within consecutive time points of the same individual— but the effect ended immediately after the isolation period when it started to increase again. External perturbations, impossible to control outside the isolation facility, modulated the salivary microbiota composition when crewmemebrs got out the isolation facility and likely started eating different types of food, getting in touch with other people, or simply visiting different places.

The finding obtained under Mars500 mission suggests that sharing the same confined environment—and possibly following the same diet regime with few variation—imbalanced the ecology of the salivary microbiota while reducing its complexity. Unfortunately, we do not know if this effect could—positively or negatively—affect the health of the hosts. However, a loss of strains in the microbiota corresponds to a loss of putative members that could be relevant under changing environmental conditions.^3,4,24^ Extending this concept we could suggest that a depauperate microbiota is less reactive to sudden changes in external conditions weaken the host adaptive capacity. However, between-subjects differences remained unaltered suggesting that individuals follow somewhat independent dynamics of their salivary microbiota (i.e. personalized dynamic). The same evidence indicates that crewmembers, though sharing the same environment, did not exchange their salivary microbiota, leading to hypothesize that quite stable personal salivary microbiota features are present in humans and confirming the between-subject effect. The fecal microbiota of the same crewmembers showed an increasing trend of similarity among subjects, especially in relation to a sharing of rare taxa,^17^ indicating that salivary microbiota has a more pronounced personalization than fecal microbiota. Indeed, from our data there are no evidences of an ecological succession, as those shown in the fecal microbiota of the same crewmembers. In support of a more pronounced personalized dynamics of salivary microbiota than of fecal microbiota in Mars500 mission, is the finding that the ASVs contributing to the temporal dynamics in the subjects were highly variable (from 3 to 28) indicating that each crewmember salivary microbiota changes were driven by a peculiar (viz. personal) set of microbial taxa.

Even if factors such as diet and time influenced the salivary microbiota composition of crewmembers, most details of the Mars500 missions are unknown. The mission was a military experiment and several outcomes are still sealed. The composition of diets for example is unknown and thus any speculation on the effect of particular food intake would not be grounded. Despite these limitations the use of 16S rRNA gene amplicon sequencing, allowed us to detect key features of human salivary microbiota under a condition that is almost impossible to replicate. The complete isolation of the participants of the mission made possible the first observation of salivary microbiota composition minimizing the effect of external perturbations.

In conclusion, the reported longitudinal analysis of human salivary microbiota confirmed the stability of the microbiota over time and suggested the presence of resilient personalized taxonomic features, which may deserve further attention. This study allowed clearly to determine the contribution of a stable and confined environment, as that of Mars500 mission, in reducing the microbiota diversity and to show the effect of a controlled diet on salivary microbiota.

## Methods

### The Mars500 experiment

Mars500 mission was conducted in 2010-2011 by Russia’s Institute of Biomedical Problems (IBMP), with extensive participation by the European Space Agency (ESA) as part of the European Programme for Life and Physical Sciences (ELIPS) to prepare for future human missions to the Moon and Mars. The whole project consisted of three isolation studies: a 14-day pilot study to test facilities and procedures used during the simulation, a 105-day pilot study involving six crewmembers, and a 520-day study that simulated a complete space flight to Mars and back. Mars500 crew was composed of six male volunteers. All crewmembers were confined in the same living space from the 3rd of June 2010 till the 4 of November 2011 when they finally stepped out of the isolation facility to come back to their normal activities. During the mission the crew was hermetically isolated from the rest of the IBMP facility. Crewmember received three type of diets, a so called “first variant” (FV), “third variant” (TV) and, after the experiment, returned to a normal (non supervised) diet regime (NR). Detailed information on the experiment are reported in Supplemental Information file. Further details are also reported in the companion Mars500 microbiology paper.^17^

### Collecting salivary samples

Saliva samples were collected individually, based on the scientific protocol and pre-confinement training, with 5ml sterile vials (Nalgene V5257-250EA). Upon completion of the saliva sampling all samples from one sampling event were put into the hatch. After that, they were removed by the responsible person of the IBMP and stored at −80°C. After being stored at −80°C in the laboratories of the IBMP for periods of at least 4 days up to 6 months, the samples were sent via World Courier. Shipping from Moscow to the University of Tuscia, Viterbo – Italy. The shipment was performed in three batches on dry ice to avoid repeated freeze-thaw cycles which lead to reduction of microbial viability. Upon arrival, samples were kept at −80°C until processing. Salivary samples were collected by crewmembers during and after the permanence in the isolation facility (Table 1): 42 samples were collected during the first simulated journey from Earth to Mars (seven time-points), 30 samples were collected during the simulated trip back home from Mars to Earth (five time-points), and other 16 samples were collected when crewmembers came back to their normal activities and were followed for additional 200 days (three time-points). Unfortunately, two samples collected in the latter stage gave no good quality DNA and were thus discharged. Crewmembers did not collect salivary samples during the simulated landing on Mars where three of them—which simulated the landing on a separate module—used a different food variant. For this reason the second food variant used was not reported in the work passing from the first food variant (FV) directly to the third food variant (TV).

**Table 1:** Number of salivary samples collected during the study.

| Subject | Earth to Mars (First diet) | Mars to Earth (Third diet) | Follow-up (No diet) | Total |
| --- | --- | --- | --- | --- |
| 5001 | 7 | 5 | 3 | 15 |
| 5002 | 7 | 5 | 3 | 15 |
| 5003 | 7 | 5 | 3 | 15 |
| 5004 | 7 | 5 | 3 | 15 |
| 5005 | 7 | 5 | 2 | 14 |
| 5006 | 7 | 5 | 2 | 14 |
| Total | 42 | 30 | 16 | 88 |

The number of samples collected during each step of the study was reported for each crewmember. Marginal totals were added for subjects and simulated journeys together with the grand total that was reported in the bottom right corner of the table.

**Table 2:** Temporal changes of salivary microbiota.

| Days | *b* | SE | *t* | *df* | *p* |
| --- | --- | --- | --- | --- | --- |
| Within-subject |  |  |  |  |  |
| 1 - 123 | 0.00103 | 0.00027 | 3.86 | 12.00 | .0023 |
| 124 - 480 | -0.00064 | 0.00012 | -5.19 | 36.00 | < .0001 |
| 481 - 720 | 0.00123 | 0.00020 | 6.16 | 14.23 | < .0001 |
| Between-subjects |  |  |  |  |  |
| 1 - 720 | -0.00004 | 0.00001 | -3.89 | 424.19 | .0001 |

Within-sample diversity was divided into three segments following change-point analysis whereas between-samples diversity was modeled on the full time period since no change-points were detected. Results of mixed effect models fitted for each segment were reported in the table. *b*, regression parameter (slope of the model); SE, standard error; *t*, t-value (also known as “standardized” regression parameter); *df*, degrees of freedom; *p*, p-value.

### Sequencing of salivary samples

DNA was extracted from salivary samples stored at −80°C using a conventional bead-beating protocol (DNeasy PowerSoil Kit, Mobio). After fluorimetry quantification (Qubit), 20 ng of environmental DNA were used as template for amplification of 16S rRNA gene using V3-V4 primers (341F and 785R) as previously reported.^25,26^ Libraries were constructed and sequenced on a MiSeq apparatus^27^ (Illumina) by BMR Genomics (Padua, Italy). Sequences are deposited at ENA database under the accession ERP119217.

### Amplicon sequence variant reconstruction

The DADA2 pipeline (version 1.14.1) was used to reconstruct amplicon sequence variants (ASVs) from illumina reads 28. Both ASV reconstruction and statistical analyses were performed in the R environment version 3.4.3 (http://www.R-project.org). For a complete description of all the step performed see Supplementary material section. Briefly, primers used for V3-V4 amplification were detected and removed using cutadapt version 1.15.^29^ Low quality reads were discarded using the `filterAndTrim` function with an expected error threshold of two for both forward and reverse read pairs. Denoising was performed using the `dada` function after error rate modelling (`learnErrors` function). Denoised reads were then merged discarding those with any mismatches and/or an overlap length shorter than 20bp (‘mergePairs’ function). Chimeric sequences were removed using the `removeBimeraDenovo` function. Taxonomical classification was performed using DECIPHER package version 2.14.0 against the latest version of the pre-formatted Silva small-subunit reference database (SSU version 132 available at: http://www2.decipher.codes/Downloads.html).^30,31^ All sequences classified as chloroplasts, mitochondria, Archaea and Eukarya were removed. A summary of retained reads in each step is reported in Table S1 and in Figure S1.

### Diversity estimation

Bacterial diversity in each sample was computed using inverse Simpson index as implemented in the diversity function of vegan package. Differences according to crewmembers, permanence in the isolation facility, and food variants were inspected using one-way analysis of variance (ANOVA). The effect of time was modeled using linear mixed models with fixed slope and random intercept. Since alpha diversity was measured multiple times on the same statistical units, crewmembers were used as random intercept factor.

Diversity across samples was inspected using different approaches. Qualitative and quantitative indexes were used to infer pairwise distances between samples. Qualitative indexes are binary indexes which take into account presence/absence of species to compute distances between samples whereas quantitative indexes are mainly based on the abundance of species.^32^ Sorensen index^33^ and un-weighted UniFrac distance^34^ were used as qualitative indexes whereas Bray-Curtis dissimilarity^35^ and weighted UniFrac distance^34^ were used as quantitative indexes. UniFrac distances were computed using the `distance` function of the phyloseq R package version 1.30.0^36^ whereas Sorensen and Bray-Curtis dissimilarity indexes were computed using the `vegdist` function of the R package vegan verison 2.5-6.^37^ Differences between salivary microbiota composition of the same crewmember at consecutive time-points (within-subject diversity) were computed using the `TBI` function of the adespatial R package verison 0.3-8.^38^ Packages `vegan` and `adespatial` use the same definition of Sorensen and Bray-Curtis distance.

Sorensen index is defined as:

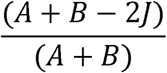

where A and B are the numbers of ASVs on compared samples, and J is the number of the ASVs shared by both samples.

Bray-Curtis index is defined as:

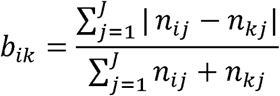

where *b*_*ik*_ is the Bray-Curtis distance between sample *i* and *k* with *J* number of species, *n*_*ij*_ is the abundance of species *j* in sample *i*, and *n*_*kj*_ is the abundance of species *j* in sample *k*^39^. The `adespatial` package computes the index by splitting the total diversity into 3 components: A (the unscaled similarity between two samples), B (the species loss between two samples), and C (the species gain between two samples). The computation is then performed using the formula (*B* + *C*)/(*2A* + *B + C*) giving the same results as the formalization given above.

Qualitative indexes rely on the assumption that all taxa are equally contributing to bacterial diversity independently from their abundance. For this reason, even extremely rare taxa may be relevant in shaping sample distribution. To relax this assumption ASVs detected in less than 5% of samples (4 samples overall) with an abundance lower than 10 were filtered out before diversity calculation.

Distances across samples were reported using non-metric multidimensional scaling (nMDS) as implemented in the `metaMDS` function of the vegan package, with 300 random starts and monotone regression.^40,41^ To test the effect of food variants, crewmembers, and time in shaping the salivary microbiota we used permutational multivariate analysis of variance on distance matrices obtained above (`adonis2` function of the vegan package with 1,000 permutations). The proportion of sum of squares from the total (namely the *R*^2^ value of permutational analysis) was used to report the percentage of variance explained by each factor included in the analysis. Before testing for differences in bacterial composition among groups is advisable to make sure that groups are homogeneously dispersed, otherwise permutational tests (such as `adonis`) may report significant results entirely due to uneven dispersion. To distinguish between actual differences in composition or differences due to dispersion we used the `betadisper` and `anova` functions (`vegan` package). P-values obtained were corrected using Benjamini & Hochberg correction (also known as false discovery rate)^42^. To avoid possible biases induced by uneven sequencing depths, read counts were scaled using DESeq2 before diversity calculation (`counts` function) 43. Scaled counts were additionally transformed using the square root of the Wisconsin double standardized counts (`wisconsin` function of the vegan package).

### Influence of time on bacterial diversity

Salivary microbiota may be affected by several factors. Sharing the same environment for a prolonged period of time may alter the composition of salivary microbiota at different levels. The bacterial composition may be altered within the same individual taken at consecutive time points but even between multiple individuals at each time point. To inspect both of these components, bacterial diversity within and between subjects, calculated as reported above, was modeled through time. Change-point analysis was used to identify specific time points which led to a decrease/increase of diversity. The optimal positioning and number of change-points for each crewmember was identified using a non-parametric cost function as implemented in the `cpt.np` function of the changepoint.np R package, version 1.0.1.^44^ The pruned exact linear time algorithm (PELT)^45^ was used to detect temporal changes in diversity within the same subject and between different subjects. The PELT algorithm searches for an optimal solution by minimizing the cost of different segmentation. We used the modified Bayes information criterion penalty term (MBIC)^46^ as penalty function for cost minimization by the algorithm. Since PELT algorithm is exact, a solution is always found for each time series so, to avoid inflation of change-points due to the presence of data coming from six different subjects, a genetic algorithm was used to fine-tune the analysis. All change-points detected were used as starting point of the genetic algorithm and a fitness function was defined as:

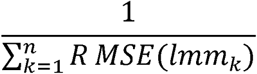

where, *RMSE*(*lmm*_*k*_) stands form the root mean square error of the generalized linear model constructed on the *k* segment and *n* is the number of segments defined by change-point analysis. High error corresponds to a low fitness value whereas low error corresponds to a high fitness value. At each step of iteration, the genetic algorithm will keep segmentation that lead to linear models with a low RMSE and discard those leading to high error models. We implemented this algorithm using the R package GA^47^ verison 3.2 with a population size of 200. Generalized linear model were fitted using crewmembers as random intercept and p-value were computed with the Satterthwaite’s degrees of freedom method as implemented in the package lmerTest^48,49^ version 3.1-1.

We assessed the persistence of the salivary microbiota across subjects using dynamic time warping algorithm implemented in the dtwclust R package (version 5.5.6)^50^. At each time point the number of subjects in which a given ASV was detected (namely the ASV’s persistence) was reported together with its abundance. Persistence matrix was scaled and centered before clustering. Centering was performed by subtracting the mean of each ASV from their persistence whereas scaling was performed by dividing each value by its standard deviation. The relation between persistence and abundance was tested by fitting a linear model (‘lm’ function of R stat package). All detail s about clustering and modelling were reported in Supplementary methods. Persistence across subjects was also used for network construction: at each time point we constructed a bipartite network by linking subjects with ASVs that were present in their salivary microbiota. Subjects were represented using squared nodes whereas ASVs were represented using round circles colored according to the groups defined above. Doing so, we generated twelve bipartite, acyclic, and undirected networks representing the salivary microbiota of all subjects at different time points. The network R package (version 1.16.0) was used for network reconstruction whereas the package ggnetwork (version 0.5.8) was used for plotting.^51,52^ The effect of time and the end of the isolation period on the number of formed/destroyed edges were tested using mixed effect models with random intercept. Subjects were taken as random intercept whereas the time and the end of the isolation period as fixed effects. P-values were computed as discussed above.

Differences along time, for each ASVs detected, were inspected by selecting drivers of within-subject beta diversity. Absolute differences between consecutive time points were fitted using linear models and the effect of time and changepoints was inspected. The slope of the models was used to assess the trend of selected ASVs in each stage. We selected ASVs showing a trend similar to what we found during changepoint detection to focus the analysis only on components of the salivary microbiota of each subject contributing to the total diversity. Finally, To inspect if “drivers of diversity” detected were more present in stable or inconsistent salivary microbiota, a hypergeometric test was performed. The test compared the the occurrence of stable ASVs (Cluster 2) in the overall population against the microbiota of single subjects seraching for significant enrichments in respect with the microbiota distribution in all subjects.^53^

## Acknowledgements

The MARS500 Programme was financed by the European Programme for Life and Physical Sciences in Space (ELIPS). The financial support of the Italian Space Agency (contract I/011/11/0) is highly remarked and acknowledged. This work was partially supported by BMR Genomics for sequencing of salivary microbiota. The authors would like to tank Professor Nicola Segata for his precious suggestions on microbiota analyses and comments on the work.

