## Supplementary materials for "Defining the resilience of the human salivary microbiota by a 520 days longitudinal study in confined environment: the Mars500 mission"

#### Mars500 mission

Mars500 mission was the first prolonged isolation experiment involving human volunteers. The experiment was conducted between 2007 and 2011 by three space agencies of different countries, namely Russia, Europe, and China. Three stages were conducted during the whole experiment with different objectives:

1. Firs stage (14-days long): testing facilities and procedures.
2. Second stage (105-days long): testing the effect of isolation on human valuteers.
3. Third stage (520-days logn): testing the effect of prolonged isolation (such as those taking plance during an hypothetical mission to planet Mars) on human valunteers.

This work focused on the last stage (the longest one) where six male volunteers were sealed inside an isolation facility built inside the Institute of Biomedical Problems (IBMP) in Moscow. Crewmembers were selected by the space agencies involved in the study with a mean age of 32 years (ranging from 27 to 38 years) and with different nationality. Three russian volunteers were selected by the Russian Federation, two European volunteers were selected by ESA, and a Chinese volunteer was selected by the Chinese National Space Administration (CNSA). During the period of isolation all conditions of a real space flight were simulated. Communications with the command centre had a delay that varies from 8 to 736 sec, reaching its peak on flight day 351. Crewmembers performed realistic activities of a long-term space mission following a weekly based program (seven days a week with two days off). Activities included physical exercises, maintenance tasks and meetings, scientific experiments, and simulated emergency events. Scientific activity was mainly focused on physiology, psychology, biochemistry, immunology, biology, and microbiology (for additional information on the experiments conducted during the mission see: <http://www.esa.int/Our_Activities/Human_and_Robotic_Exploration/Mars500/Scientific_protocols>). The full timetable of the project is reported in Table S1.

The isolation facility consisted of four interconnected modules plus one external module aimed to simulate Martian surface (“Simulator of the Martian surface”, SMS module, 1200m^3^). The total volume of the habitat modules was 550m^3^ and single modules were divided as follows:

- Medical module (EU-100, 100m^3^): two medical beds, a toilet and medical equipment for routine examinations and diagnostic investigations. The module was used for quarantine in case a crewmember became ill.
- Habitable module (EU-150, 150m^3^): the main living module. Composed of 6 bedrooms (one for each crewmember) a kitchen, a dining room, and a living room. A common toilet was also present and each bedroom was equipped with a bed, a desk, a chair, and shelves.
- Mars landing module simulator (EU-50, 50m^3^): only used during the landing simulation. The module was designed to accommodate three crewmembers so it was equipped with only 3 bunk beds. In addition the module contained: 2 work stations, a toilet, a control and data collection system, a video control and communications system, gas analysis system, air-conditioning and ventilation system, sewage system and water supply, and a fire alarm and suppression system.
- Storage module (EU-250, 250m^3^): largest habitat module, it was divided into 4 compartments. Compartment one contained a fridge for storage of food, compartment two was used for storage of non-perishable food, compartment three contained the experimental greenhouse, and compartment four was composed of a bathroom, sauna and gym.

Modules included all equipment needed for conducting the study, such as ventilation and air supply, water supply, sewage, electrical installations, air and water monitoring systems and partial recycling, medical and emergency equipment. The crew lived in those modules under controlled conditions simulating an atmospheric environment at normal barometric pressure. To assess the microbial load in the isolation facility, the gas media was sampled on a regular basis (at least once a month) together with selected surfaces from the habitat modules that were swabbed over time by the crewmembers. A previous study conducted by the German Aerospace Centre,^1^ highlighted that, despite the presence of air filters, a complete sterile environment is not attainable. The presence of complex bacterial communities able to adapt to different environmental conditions was indeed confirmed posing the accent on the need of developing effective countermeasures aimed to maintain a healthy environment during long-term missions.

Like in a real space mission, both water and food supplies were limited and their composition followed the standards reported in the International Space Station (ISS) guidelines.^2^ Water supplies were divided into two separated systems, one was used for cooking and drinking, whether the other was directly connected to the centralised system of water supply in Moscow and was mainly used for sanitary-hygienic needs. Water quality in the first system was checked every two weeks and no episodes of increased microbial contamination were reported for the whole duration of the mission. Food was rationed with a nutrient content that respected the World Health Organization (WHO) guidelines as well as the standards reported for the ISS crew. Three different food rations were provided to the crew according to the phase of the mission:

- First variant: flight from the Earth to Mars - 250-day flight, from the 1st day till the 250th of the mission
- Second variant: only to the three crewmembers that simulated the landing on Mars - 20-day simulation, from the 251th till the 270th day of the mission
- Third variant: trip back home - 250-day flight, from the 271th day till the 520th day of the mission included

#### Experimental design

Salivary samples were collected by crewmembers throughout the whole 520-day mission except during the simulated landing on Mars since the Mars landing module simulator was not equipped with sampling supplies. Crewmembers were followed for additional 200 days after they left the isolation facility and got back to their normal activities to inspect how long it would take to re-establish a baseline condition after a prolonged period of isolation. We collected a total of 88 samples from the first day of the mission till the end of the follow-up (720 days in total, Figure S1).

#### Amplicon sequence analysis

Amplicon sequences were analysed using the DADA2 package version 1.6.0^3^ in R version 3.4.3 (<http://www.R-project.org>). Primers used for PCR amplification were removed using cutadapt version 1.15 in paired-end mode.^4^ Sequences that did not contain primers were discarded and both pairs were removed if one of them (R1 or R2) did not fulfill the filtering criterion (90.52% of raw sequences were retained). To infer amplicon sequence variants (ASVs) with a modest memory requirement, the big-data approach reported at: <https://benjjneb.github.io/dada2/bigdata.html> was followed. Forward reads were trimmed at 270bp and reverse reads were trimmed at 190 bp to remove low quality segments. Reads (and their respective forward or reverse read) containing ambiguous bases and more than two expected errors were filtered out (82.05% of processed reads were retained). Errors were then modelled and corrected using the DADA2 algorithm with default parameters. The denoised output reads were merged and all reads with any mismatches and an overlap length shorter than 20bp were removed (93.26% of trimmed reads were correctly merged). Chimeric sequences were identified and removed using the removeBimeraDenovo function with the consensus method (29.57% of analysed sequence were flagged as chimeric and removed from subsequent analyses). Sequence variants were taxonomically annotated using DECIPHER package version 2.6.0 against the Silva SSU reference database version 132.^5,6^ Sequences aligning to chloroplasts, mitochondria, Archaea and Eukaryotes were removed (0.08% of processed reads and 5.16% of SVs). A summary of retained reads in each step is reported in Table S2 and in Figure S2.

#### Amplicon sequence variant evaluation

Three samples coming from three subjects were sequenced three times each to validate the workflow used for inferring sequence variants (SVs). DNA was extracted independently for each replicate and sent to an external service for PCR amplification and sequencing (see Materials and Methods section pf the main manuscript). Replicates were subjected to DADA2 pipeline together with the other samples and the accuracy of the pipeline was then evaluated on replicated samples. Two main parameters were evaluated: a. the correlation coefficient between replicates, and b. the overall accuracy. The accuracy was computed as:

$$\frac{(TP+TN)}{(TP+TN+FP+FN)}$$

where, $TP$ is the number of SVs detected in both replicates; $TN$ is the number of SVs not detected in both replicates; $FP+FN$ is the number of SVs detected in one replicate but not in the other one. Accuracy was estimated by comparing replicates in pairs (three replicates equals to three possible pairs). The non-parametric Spearman’s rank correlation coefficient (rho) was used to measure monotonic relationships between SVs in all replicates. Values of Accuracy and rho for each sample sequenced were reported in Table S3 whereas the log_2_-transformed counts of SVs in each replicate was reported in Figure S3.

Rarefaction curves were computed using the R package vegan, version 2.5.6^7^ and reported in Figure S4. Slopes were calculated as the derivative of the rarefaction curves to ensure that the sequencing effort was enough for all samples. Selected replicates were reported in Table S4 along with the number of reads assigned to SVs.

Good’s coverage estimator was calculated using the formula:

$$(1-\frac{n}{N})\cdot100$$

where, $n$ is the number of sequences found only one time in a specimen (singletons) and $N$ is the total number of sequences assigned to an OTU in that specimen.^8^ Good’s coverage estimator ranged between 99.99% and 100.00% across all samples indicating that roughly 0.01% of the reads in a sample came from ASVs that appear only once in that sample (Table S5).

#### Phylogenetic reconstruction

For phylogenetic analysis all SVs were aligned with SINA 9 using the same reference database that was used for taxonomical classification (Silva SSU reference database version 132). Since the V3-V4 region of the 16S rRNA gene has no sufficient taxonomical resolution to completely resolve bacterial taxonomy,^10–12^ SVs assigned to the same Phylum were constrained to strict monophyly whereas SVs with an unknown phylum attribution were left unconstrained. Maximum likelihood trees were then calculated with RAxML (version 8.2.9)^13^ under the GAMMA model of rate heterogeneity and the GTR substitution matrix. A phylogenetic tree was constructed with 20 maximum likelihood (ML) searches on 100 randomized stepwise addition parsimony trees (Figure S5). The tree was rooted using midpoint rooting strategy as implemented in the R package phangorn, version 2.5.5.^14^ The resulting tree was used to calculate distances between samples using UniFrac metrics^15^ (both weighted and unweighted).

#### Effect of diet on salivary microbiota

The effect of diet on the main genera detected was inspected using the Mann-Whitney test on reads counts previously scaled used DESeq2 (counts function with norm = T parameter specified) and transformed into relative abundances. We repeated the test for all contrasts of the three diets used to feed crewmebers during the mission and p-values were finally corrected using the Benjamini & Hochberg correction (p.adjust function of R). The mean abudance during each diet was reported togheter with the standard error on the mean (Figure S6).

#### Stable versus inconsistent microbiota

To detect stable and inconsistent microbiota amplicon sequence variants were clustered according to their persistence along the mission and during the follow-up. Dinamyc time warping (DTW) algorithm was used to measure similarities between temporal sequences of ASV persistance, as implemented in the dtwclust R package (version 5.5.6).^16^ Silouette score was used to inspect the distance between clusters prodeced and select the optimal number of cluster for the DTW algorithm (silhouette function of the cluster R package version 2.1.0).^17^ The optimal number of clusters was set to two (Figure S8 panel a) resulting in a sharp separation between two groups of ASVs with different resilience levels (Figure S8 panel b). The abundance of the two groups of ASVs was reported in panel c of Figure S8. The number of specimens in which a given ASV is detected (persistence) should correlate with the abundance of that ASV. The abundance of all ASVs in different sampling sites and at different time points was indeed significantly correlated with the number of subjects in which those ASVs were detected, reporting an $R^{2}=.72$ (linear regression on log-transformed abundance; $b=10.09$, 95% CI $[9.81$, $10.38]$, $t(1926)=70.27$, $p<.001$, Figure S9).

### Supplementary Tables

Table S1: Full experiment timetable. The Mars500 project consisted of three isolation experiments: two pilot suties (14 and 105 days, completed in November 2007 and July 2009, respectively) and the main 520-day study. All studies were conducted inside an isolation facility that was buit in the Institute of Biomedical Problems (IBMP) of the Russian Academy of Sciences located in Moscow.

| Date | Event |
| --- | --- |
| November 2007 | First pilot study (14-day simulation for testing facilities and operational procedures) |
| 31 March 2009 | Start of the second pilot study (105-days, four Russian and two European crewmembers) |
| 14 July 2009 | End of the second pilot study |
| 23 March 2010 | European candidates for the 520-day crew announced |
| 18 May 2010 | The whole 520-day crew announced (three Russian, two European and one Chinese) |
| 3 June 2010 | Start of the 520-day isolation study |
| 3 Jun 2010 | Start of the main experiment - hatch closed, lift off |
| 15 Jun 2010 | Undocking from orbital assembly laboratory |
| 23 Jun 2010 | Transfer to heliocentric orbit towards Mars |
| 24 Dec 2010 | Shifting to spiral orbit towards Mars |
| 1 Feb 2011 | Entering circular orbit around Mars |
| 1 Feb 2011 | Mars Lander hatch opening |
| 8 Feb 2011 | Completion of loading, Lander hatch closure |
| 12 Feb 2011 | Undocking, landing on Mars |
| 14, 18 and 22 Feb 2011 | Egresses on Martian surface |
| 23 Feb 2011 | Ascent, beginning of quarantine |
| 24 Feb 2011 | Docking with interplanetary craft |
| 26 Feb 2011 | End of quarantine |
| 27 Feb 2011 | Habitation module hatch opening |
| 27 Feb 2011 | Crew transfer to Habitation module |
| 28 Feb 2011 | Lander loading with trash ends |
| 1 Mar 2011 | Hatch closure, Lander undocking |
| 2 Mar 2011 | Entering into spiral orbit away from Mars |
| 7 Apr 2011 | Transfer to heliocentric orbit towards Earth |
| 15 Sep 2011 | End of communications delay, switchover to voice communications |
| 13 Oct 2011 | Shifting to spiral orbit towards Earth |
| 4 Nov 2011 | End of 520-day study, crew landing on Earth |

Table S2: Number of reads retained during the amplicon sequence variant inference. The number of reads retained fater each step of analysis is reported in different columns: sample_id, id of the sample included in the study; raw, sequencing depth in the raw file; no.adapt, number of reads retained after primer removal step; filtered, number of quality filtered reads; merged, number of forward and reverse reads correctly merged; no.chim, number of reads retained after chimeric removal step; bacteria, read assigned (at least) to Bacteria domain; no_chloroplast, number of reads retained after Chloroplast removal; no_mitochondria, number of reads retained after Mitochondria removal.

Table S3: Accuracy and correlation between replicates. Contrasts are reported in the contrast column whereas subject id refers to the subject that has been processed. Spearman’s rank correlation coefficient was reported in the “rho” column. Accuracy was estimated as reported in Supplementary method section.

| Subject id | Contrast | rho | Accuracy |
| --- | --- | --- | --- |
| 5001 | a vs. b | 0.98 | 0.97 |
| 5001 | a vs. c | 0.97 | 0.97 |
| 5001 | b vs. c | 0.98 | 0.98 |
| 5003 | a vs. b | 0.98 | 0.98 |
| 5003 | a vs. c | 0.94 | 0.98 |
| 5003 | b vs. c | 0.95 | 0.98 |
| 5005 | a vs. b | 0.97 | 0.96 |
| 5005 | a vs. c | 0.98 | 0.96 |
| 5005 | b vs. c | 0.97 | 0.96 |

Table S4: Number of reads assigned to SVs for each replicate. Replicate with the highest number of sequences assigned to SVs were selected and retained for subsequent analyses whereas other replicates were removed. Selected samples were marked with an asterisk.

| Sample_id | subject_id | replicate | n |
| --- | --- | --- | --- |
| 79529 | 5001 | a | 51919 |
| 79517* | 5001 | b | 76972 |
| 79518 | 5001 | c | 74631 |
| 79471 | 5003 | a | 54619 |
| 79519* | 5003 | b | 62802 |
| 79520 | 5003 | c | 60465 |
| 79485 | 5005 | a | 31151 |
| 79521 | 5005 | b | 37392 |
| 79522* | 5005 | c | 51345 |

Table S5: Good’s coverage estimator for each sample. Sample_id, id of samples included in the study; Clones, number of reads correctly processed and assigned to an ASV; ASVs, number of sequence variants detected for each sample; Singletons, number of ASVs with only one reads mapped in a sample; Doubletons, number of ASVs with two reads mapped in a sample; Good_coverage_estimator, Good’s coverage estimator.

Table S6: Overall abundance and percentage of reads assigned to bacterial Phyla. The abundace column reports the number of reads assigned to all ASVs with the same phylum classification. The number of ASVs assigned to each phylum was reported in the N column. The baundance was divided by the total amount of reads produced for the whole experiment and reported as percantage. Bacterial ASVs with no phylum classification were reported as “Unknown”.

| phylum | Abundance (reads’ count) | Percentage | N |
| --- | --- | --- | --- |
| Firmicutes | 2101490 | 48.45 | 516 |
| Bacteroidetes | 1050115 | 24.21 | 526 |
| Actinobacteria | 590589 | 13.62 | 205 |
| Proteobacteria | 330630 | 7.62 | 177 |
| Fusobacteria | 161541 | 3.72 | 136 |
| Patescibacteria | 81283 | 1.87 | 52 |
| Epsilonbacteraeota | 11736 | 0.27 | 14 |
| Spirochaetes | 4219 | 0.10 | 42 |
| Tenericutes | 1118 | 0.03 | 9 |
| Synergistetes | 276 | 0.01 | 4 |
| Unknown | 4543 | 0.10 | 247 |

Table S7: Analysis of variance model. The effect of diets, subjects, and the permanence into the isolation facility (experiment) on alpha diversity (inverse-Simpson index) was explored using a one-way analysis of variance (ANOVA). No significant effect was found. F, value of F-statistic; df1 and df2, degrees of freedom available for the considered factor and total degrees of freedom; MSE, mean square error; p, p-values corresponding to the given F-statistic; ges, effect-size measured as generalized eta-squared.

| Effect | F | df1 | df2 | MSE | p | ges |
| --- | --- | --- | --- | --- | --- | --- |
| Experiment | 2.95 | 1 | 84 | 84.76 | .090 | .034 |
| Food | 0.98 | 1 | 84 | 84.76 | .325 | .012 |
| Subject id | 0.30 | 1 | 84 | 84.76 | .585 | .004 |

Table S8: Mixed effect model fitting with Satterthwaite approximation. The effect of days on the overall bacterial diversity (inverse-Simpson index) was explored through a mixed effect model with random intercept. Subjects were taken as blocking factor and p-values were obtained using the Satterthwaite approximation implemented in the lmerTest R package (version 3.1). No significant effect was found. $b$, regression parameter (Days: slope of the model; Post-isolation: difference in the intercept); SE, standard error; $t$, t-value (also known as “standardized” regression parameter); $df$, degrees of freedom; $p$, p-value.

| Effect | $b$ | SE | $df$ | $t$ | $p$ |
| --- | --- | --- | --- | --- | --- |
| Days | 0.002 | 0.004 | 82.03 | 0.47 | 0.641 |

Table S9: Mixed effect model with Satterthwaite approximation. The effect of time on the number of formed and destroyed edges in the community network was tested using mixed effect models. Subjects were used as blocking factor and p-values were obtained using the Satterthwaite approximation implemented in the lmerTest R package (version 3.1). $b$, regression parameter (Days: slope of the model; Post-isolation: difference in the intercept); SE, standard error; $t$, t-value (also known as “standardized” regression parameter); $df$, degrees of freedom; $p$, p-value.

| Effect | $b$ | SE | $df$ | $t$ | $p$ |
| --- | --- | --- | --- | --- | --- |
| Formed |  |  |  |  |  |
| Days | -0.05 | 0.02 | 74.88 | -3.19 | 0.002 |
| Post-isolation | 28.19 | 10.06 | 75.71 | 2.80 | 0.006 |
| Destroyed |  |  |  |  |  |
| Days | -0.04 | 0.02 | 74.94 | -2.32 | 0.023 |
| Post-isolation | 13.59 | 11.61 | 75.75 | 1.17 | 0.245 |

Table S10: Number of ASVs reporting a trend of diversity similar to the whole within-subject diversity. Results of the linear models were reported for each ASVs and for each subject. Subject, the id of the crewmember; ASV, the id of the amplicon sequence variant detected; F, value of F statistic; df1 and df2, degrees of freedom used for F staistic calculation; p, p-values ; p.adj, Benjamini & Hochberg corrected p-values; domain, phylum, class, order, family, and genus, taxonomic classification of ASVs; cluster, the cluster to which a given ASV belongs.

### Supplementary Figures

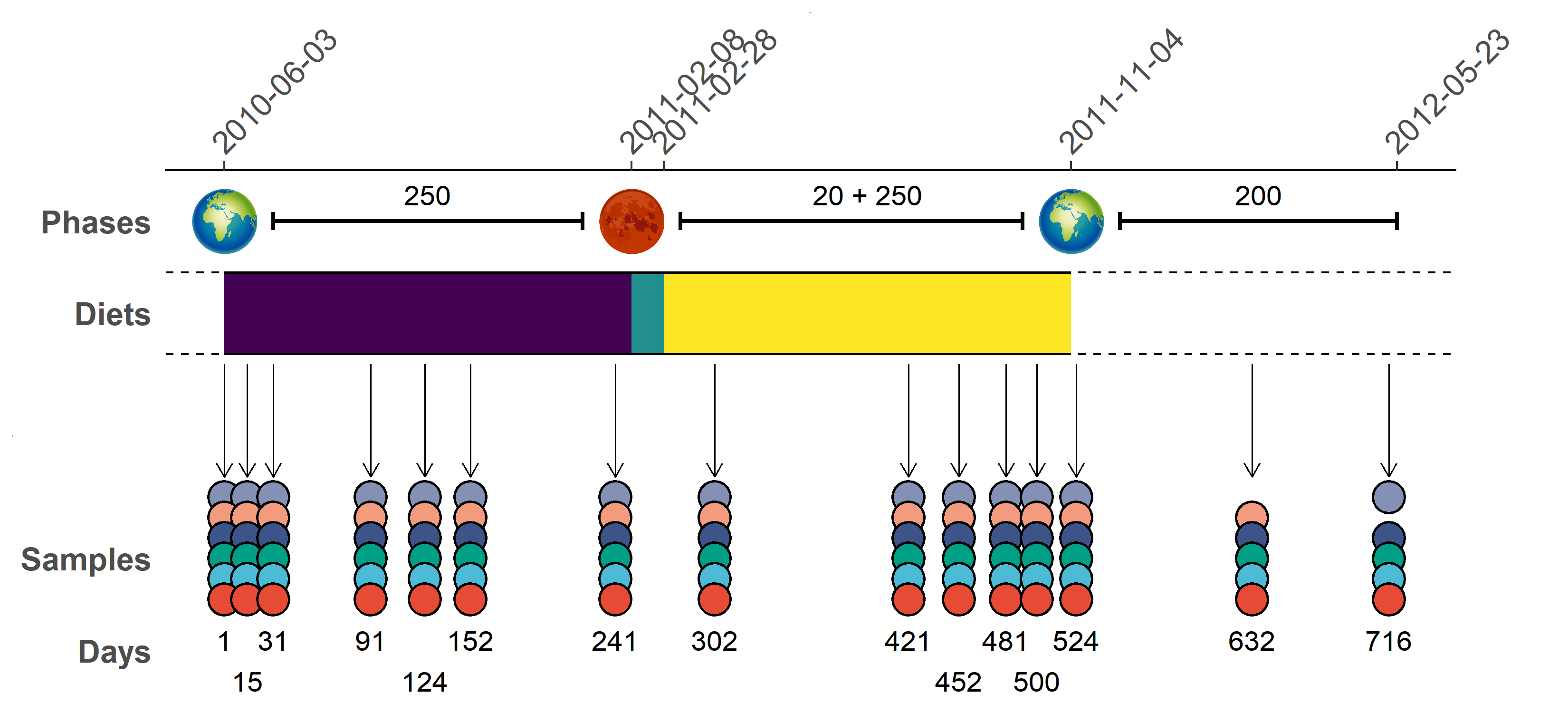

Figure S1: Timeline of Mars500 mission. The phases of the mission were reported in the top part of the plot togheter with their length in days. The second phase included the landing simulation (20 days) as well as the trip back to Earth (250 days). Diets supplied to crewmembers were reported using different colors, violet for the first food variant, cyan for the second food variant, and yellow for the third food variant. Samples were reported using one point for each crewmember (subject 5001 red, 5002 pale blue, 5003 green, 5004 dark blue, 5005 ochre, and 5006 gray). The days of isolation were reported in the bottom part of the plot.

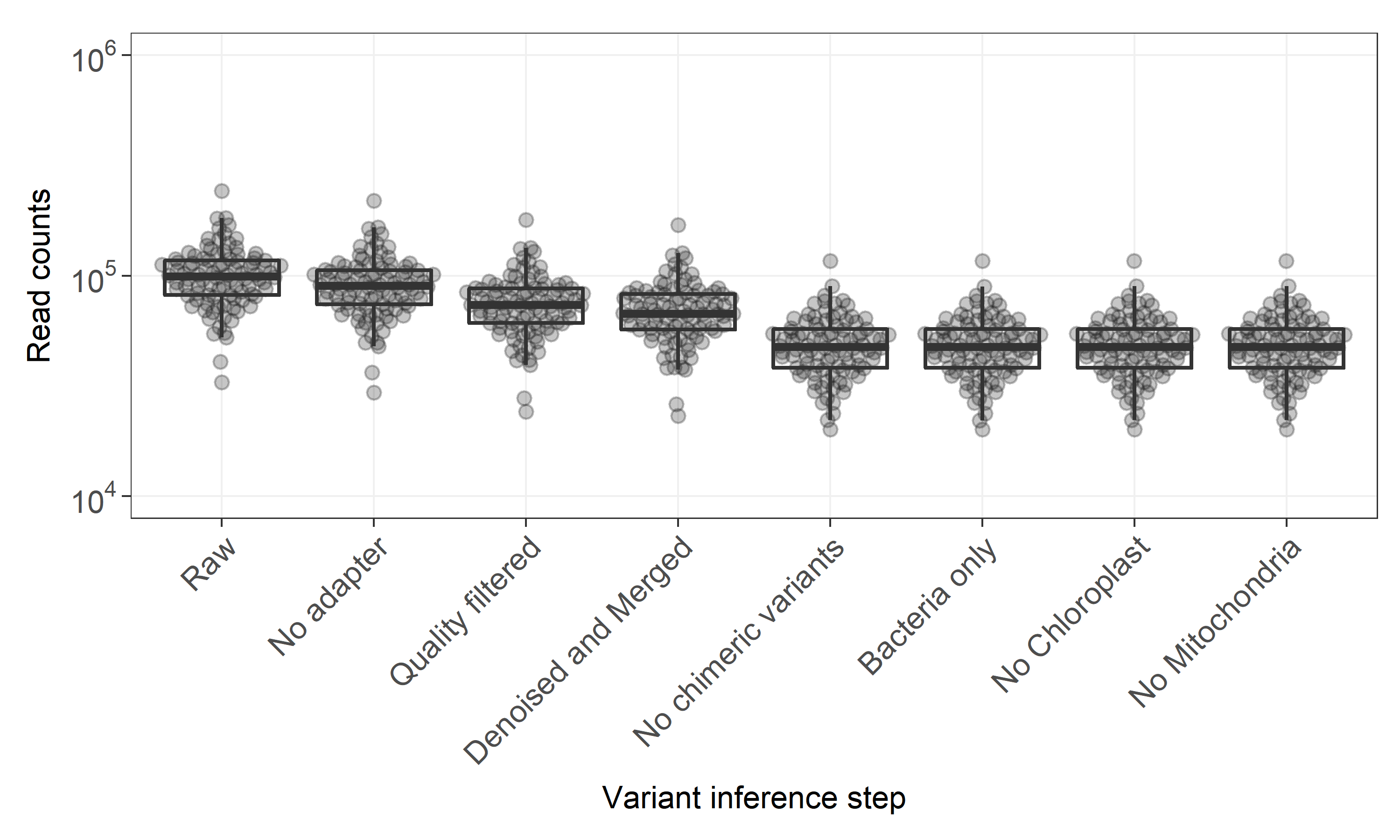

Figure S2: Number of reads retained after each step of the DADA2 workflow.

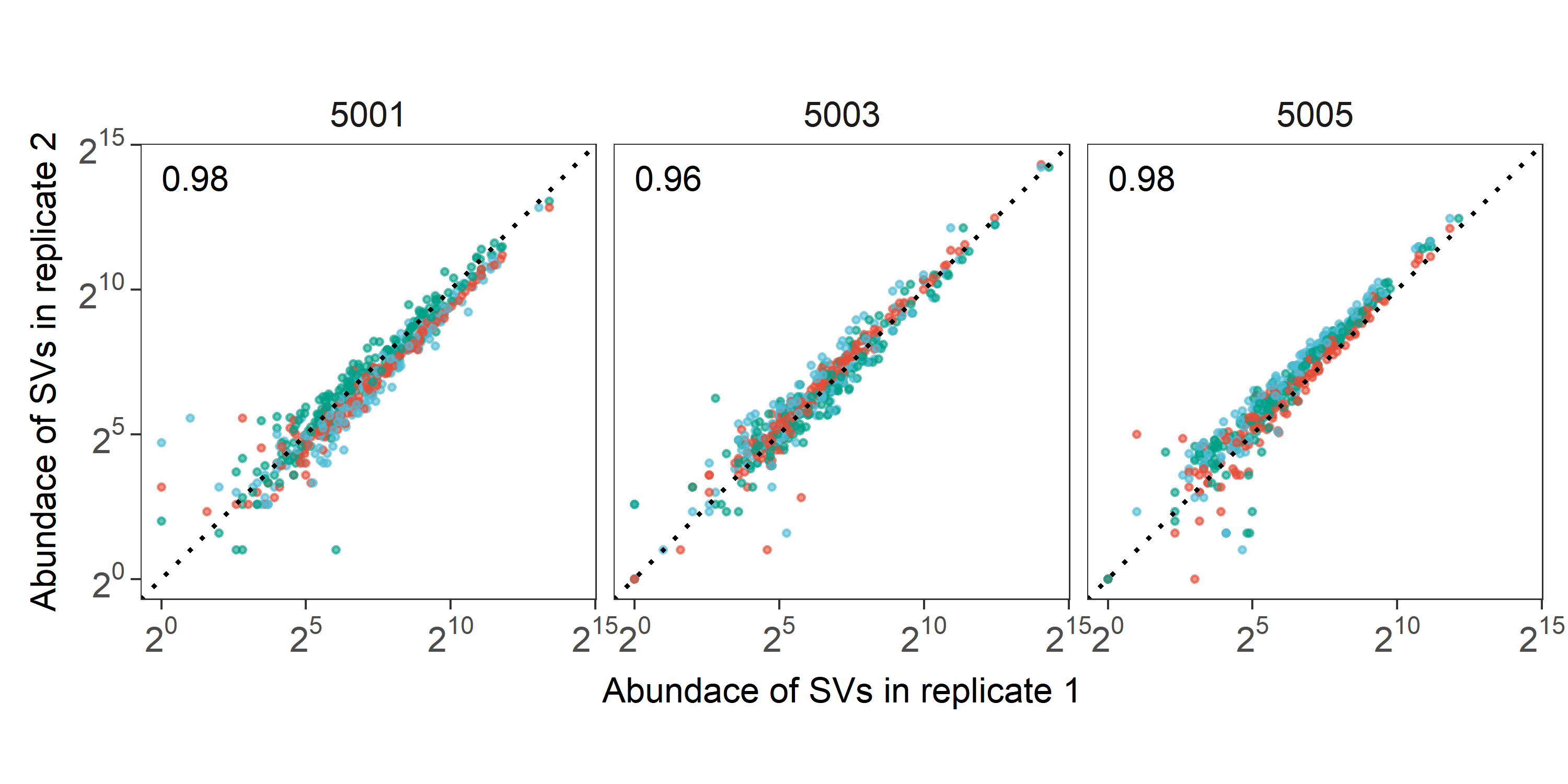

Figure S3: Counts of individual SVs for each replicate. The abundance of every SV is reported for the three pairwise combinations using the log2-scale. Colors represent comparisions between replicates (a vs. b red dots, a vs. c blue dots, and b vs. c green dots) whereas subjects were reprted in different panels. Dotted lines represent a perfect correlation (namely a line with slope equal to one and intercept equal to zero) whereas the average value of the Spearman’s correlation coefficent was reported at the top left of each panel.

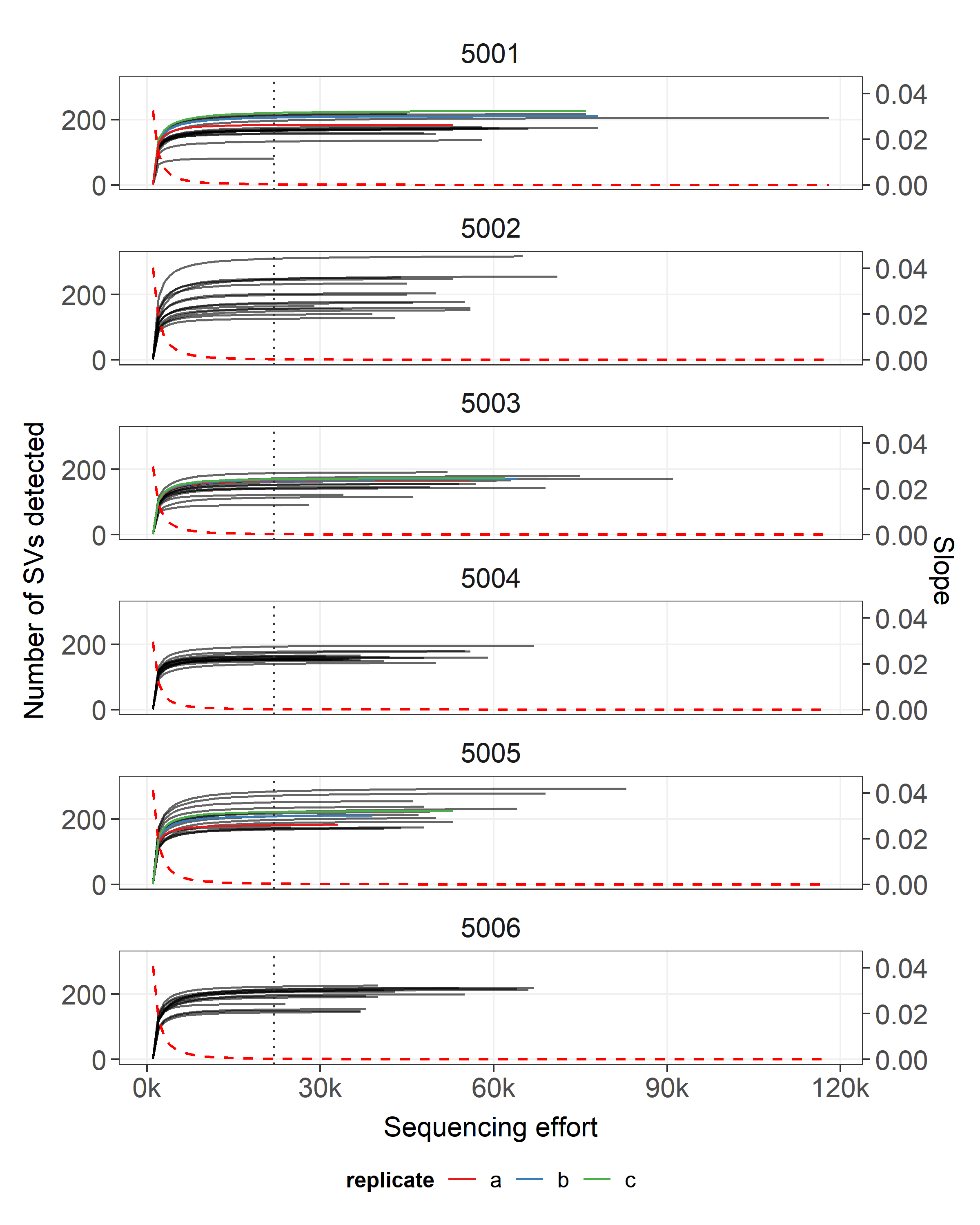

Figure S4: Number of SVs detected (left side vertical axis) with increasing sequencing effort (horizontal axis). The average slope of rarefaction curves was reported with a dashed line (right side vertical axis). Vertical lines represent the sample with the lowest number of reads assigned to a SVs. Rarefaction curves were computed with an increasing step of 1000 reads. Samples that were sequenced multiple times (tecnical replicates) were reported with different colors.

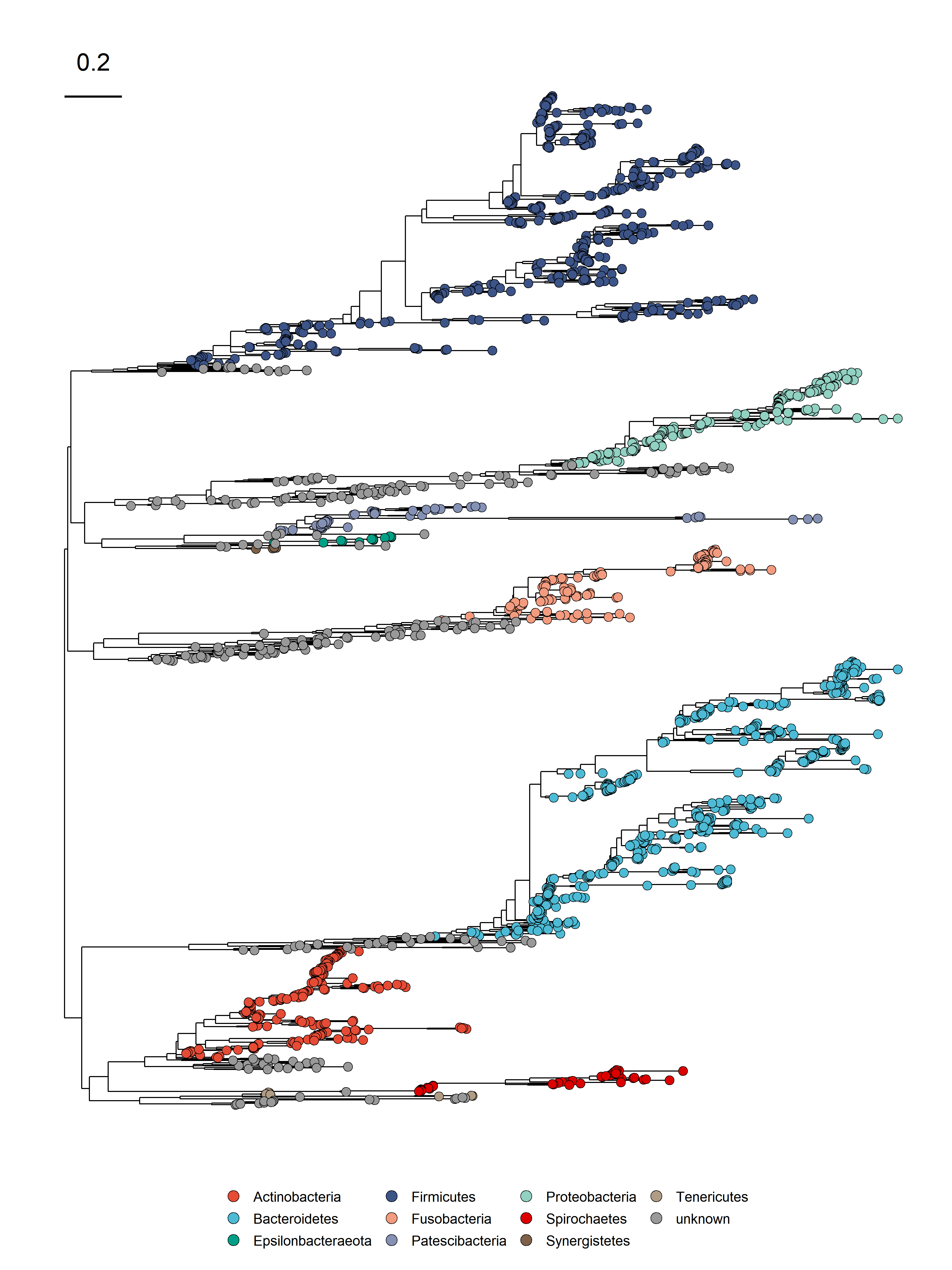

Figure S5: Phylogenetic reconstruction of ASVs. Phyla were reported with different colors whereas SVs with unknown phylum attribution were reported in gray. The distance scale is reported in the top left of the plot.

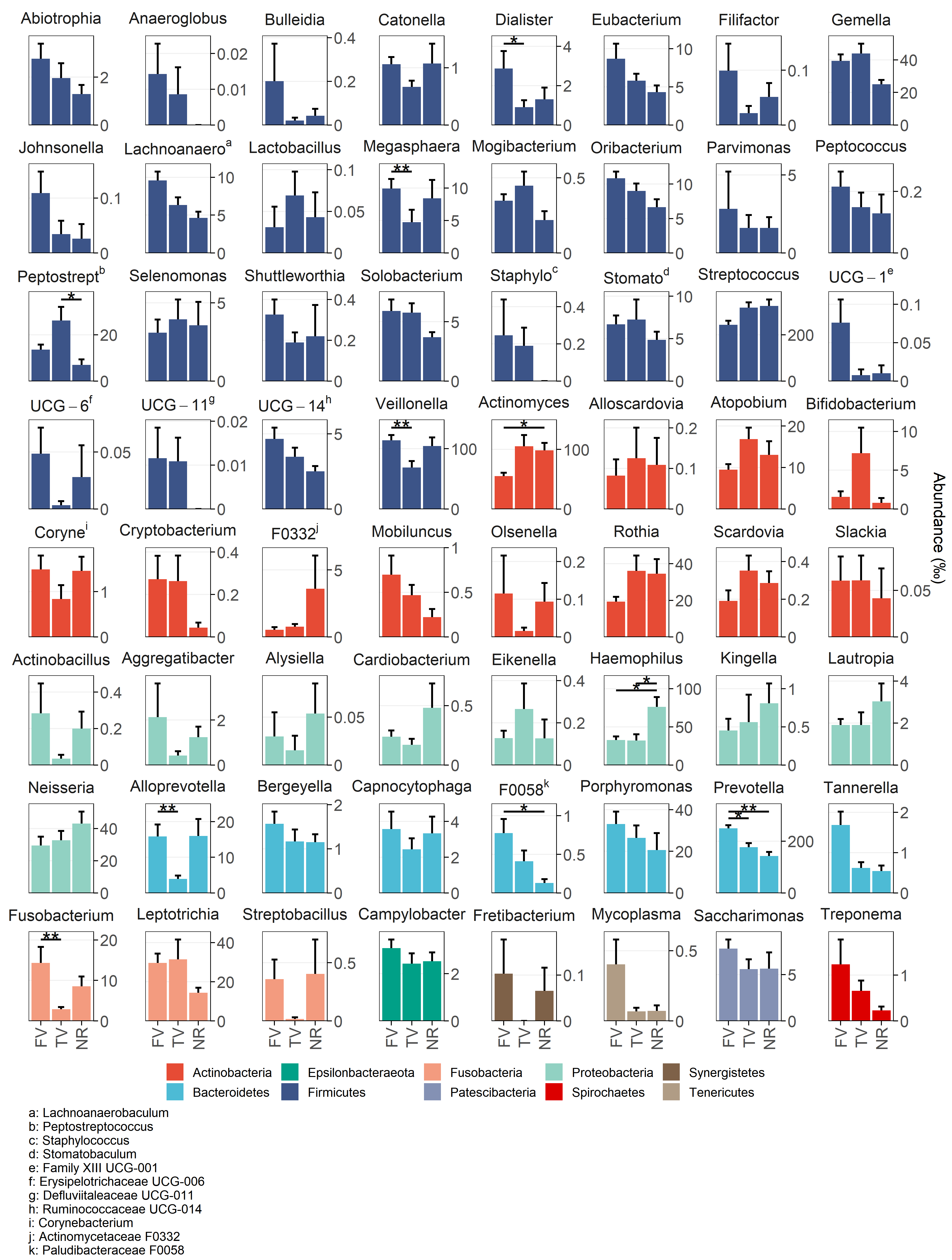

Figure S6: Bactrerial genera detected in different diets. Normalized counts were transformed into parts per thousand and reported in different panels according the the genus attribution. Diets are reported in the x-axis whereas the mean abundace was reported in the y-axis togheter with the standard error on the mean (error bars). Mann-Whitney test detected significant chenges in bacterial distribution according to diet that were reported using asterisks (*, p < 0.05; **, p < 0.01; Benjamini & Hochberg correction).

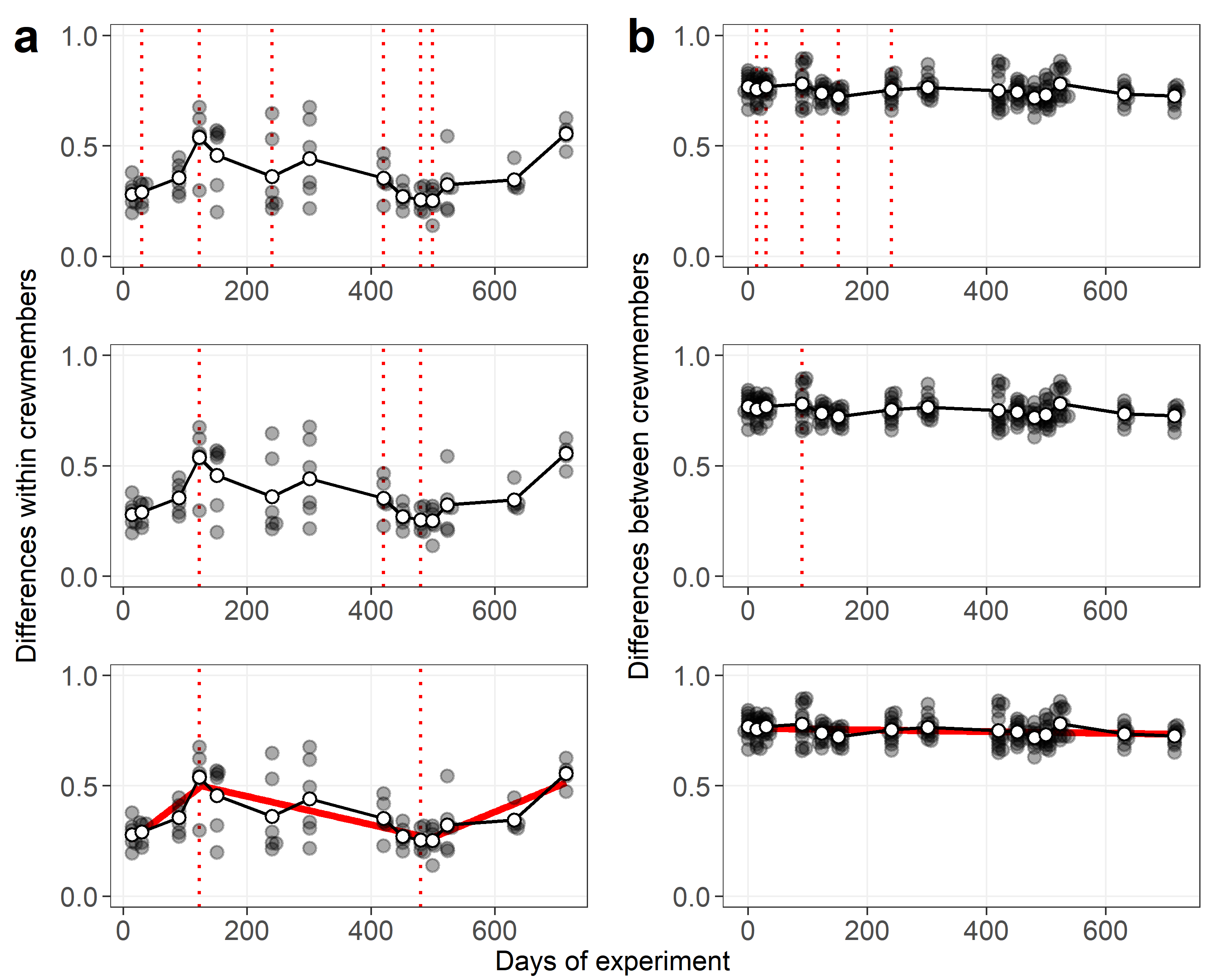

Figure S7: Changepoint detection analysis based on differences within crewmembers (a) and differences between crewmembers (b). Each step of analysis is reported in different panels from top to bottom: changepoint analysis on all crewmembers, genetic algoritm selection of changepoints producing relevant trends of beta diversity, linear model/s fitting after removal of non-significant changepoint/s.

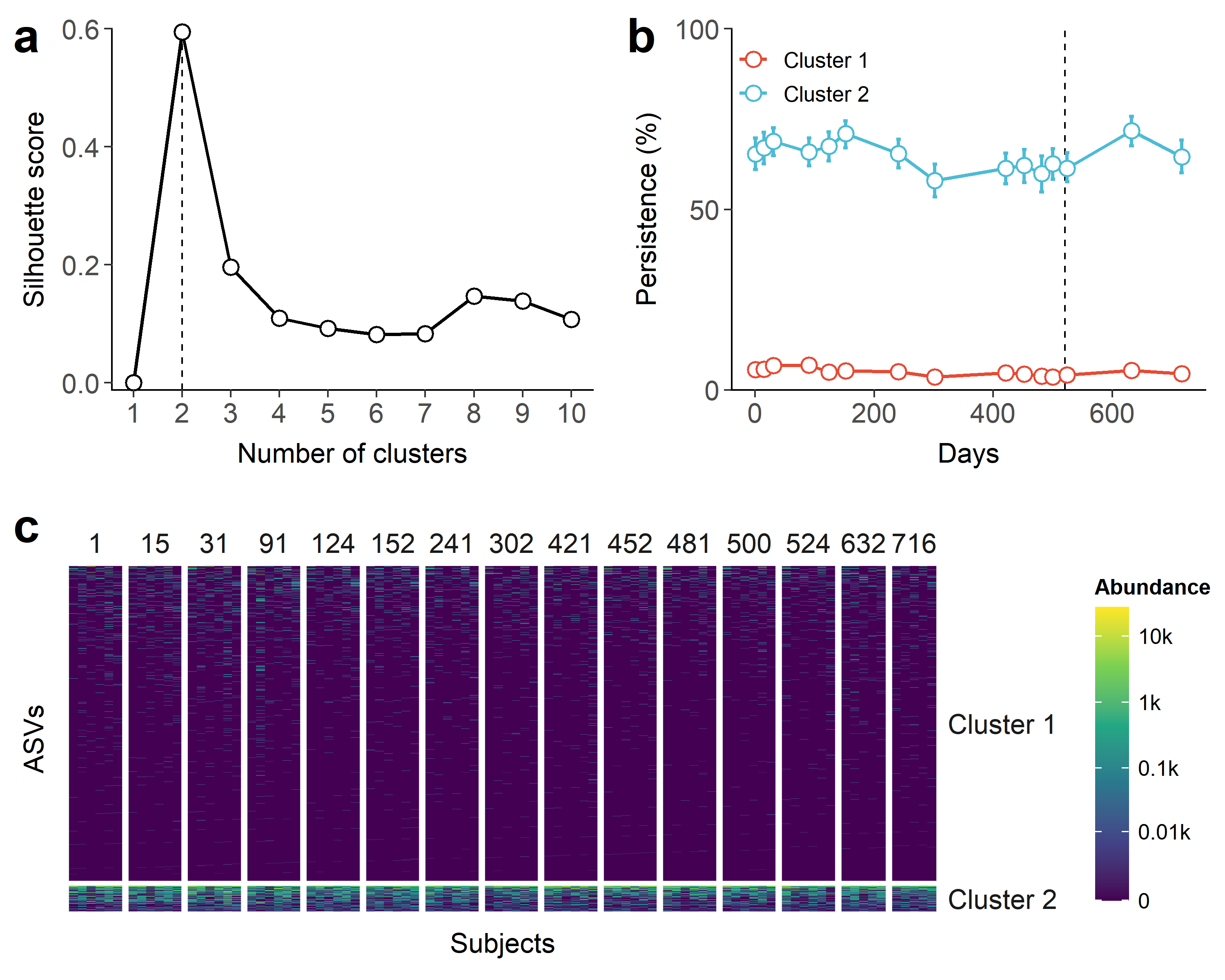

Figure S8: Clustering of ASVs based on time-resolved persistence The number of subject in which a given ASV was detetcted was computed at each time point (persistence). The dynamic time warping (DTW) approach was used to cluster ASVs with increasing number of clusters. a) The number of clusters maximizing the Silhouette score was chosen as the best value. b) The prevalence of ASVs assigned to each cluster was plotted along time, whereas c) abundance values was reported as a heatmap.

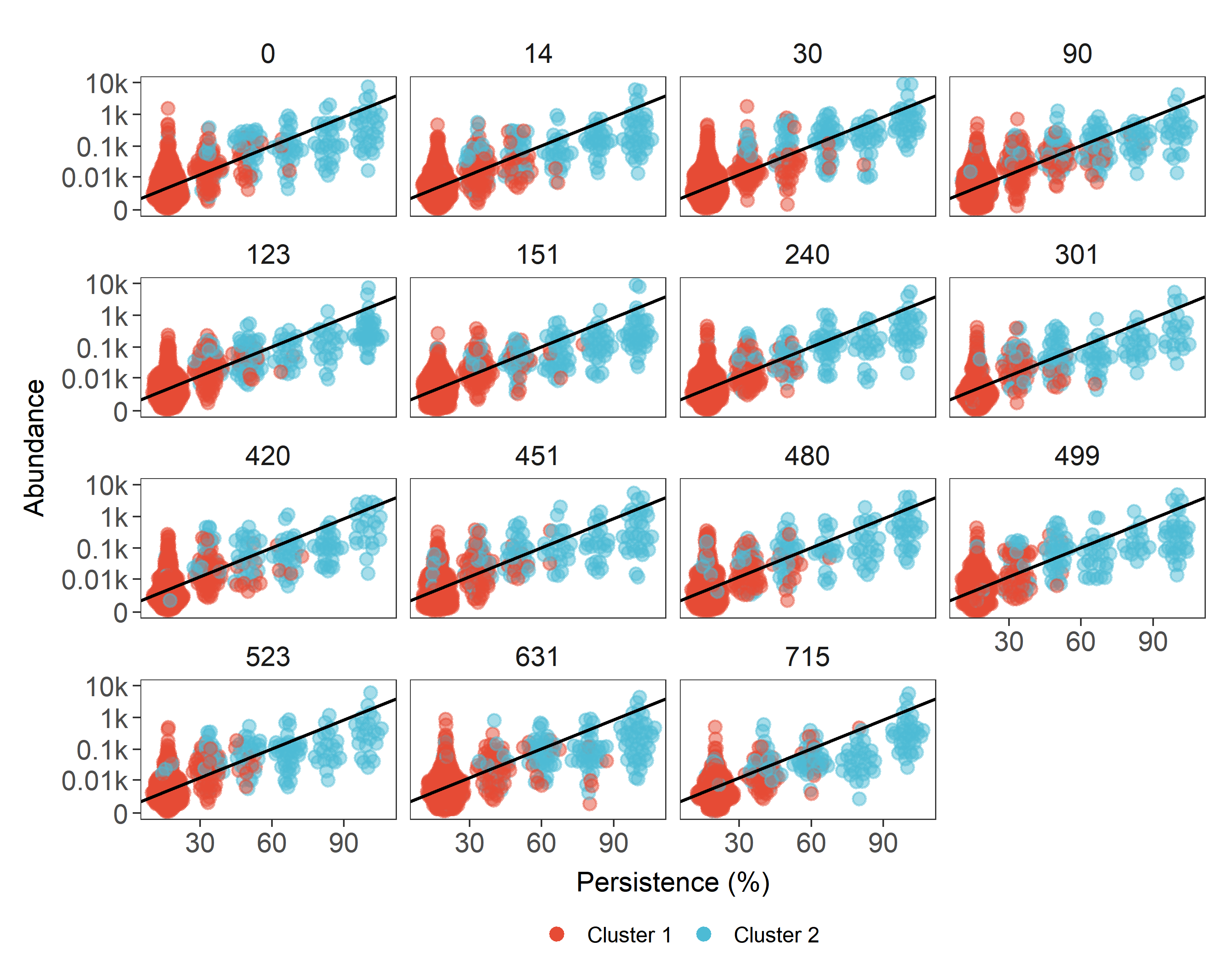

Figure S9: Persistence and abundance of ASVs detected at different time points. Persistence was expressed as the number of samples in which a given ASV was detected whereas abundance was expressed as log-normalized number of reads assigned to a given ASV (linear regression was reported as a black line (95% CI [9.81, 10.38], t(1926) = 70.27, p < 0.001). Clusters were reported using different colors.
